# Multi-region calcium imaging in freely behaving mice with ultra-compact head-mounted fluorescence microscopes

**DOI:** 10.1101/2023.10.30.564709

**Authors:** Feng Xue, Fei Li, Ke-ming Zhang, Lufeng Ding, Yang Wang, Xingtao Zhao, Fang Xu, Danke Zhang, Mingzhai Sun, Pak-Ming Lau, Qingyuan Zhu, Pengcheng Zhou, Guo-Qiang Bi

**Author notes:** These authors contributed equally to this work (Feng Xue, Fei Li, Ke-ming Zhang).

## Abstract

To investigate the circuit-level neural mechanisms of behavior, simultaneous imaging of neuronal activity in multiple cortical and subcortical regions is highly desired. Miniature head-mounted microscopes offer the capability of calcium imaging in freely behaving animals. However, implanting multiple microscopes on a mouse brain remains challenging due to space constraints and the cumbersome weight of equipment. Here, we present TINIscope, a Tightly Integrated Neuronal Imaging microscope optimized for electronic and opto-mechanical design. With its compact and lightweight design of 0.43 g, TINIscope enables unprecedented simultaneous imaging of behavior-relevant activity in up to four brain regions in mice. Proof-of-concept experiments with TINIscope recorded over 1000 neurons in four hippocampal subregions and revealed concurrent activity patterns spanning across these regions. Moreover, we explored potential multi-modal experimental designs by integrating additional modules for optogenetics, electrical stimulation or local field potential recordings. Overall, TINIscope represents a timely and indispensable tool for studying the brain-wide interregional coordination that underlies unrestrained behaviors.

## INTRODUCTION

A fundamental goal of neuroscience is to understand the neural basis of behavior and cognition, which involve the coordinated activity of multiple brain structures distributed across cortical and subcortical areas[1-3]. Several studies have leveraged multi-site electrophysiological recordings to reveal the brain-wide circuit-level interactions underlying complex brain functions[4-7]. Compared to electrophysiological approaches, calcium imaging has indispensable advantages in research requiring cell-type specificities, precise spatial positions, or long-term tracking of individual neurons. However, due to technical constraints of imaging systems, most multi-region calcium imaging data were collected in the superficial brain areas of head-stabilized animals[8-10], limiting the applicability of certain behavior assays or the exploration of targeted deep brain regions. Thus, it is highly desirable to have techniques capable of imaging multiple cortical and subcortical regions spanning different depths (i.e., thalamus, hippocampus, and cortex) simultaneously during unrestrained behaviors, especially in mice[11].

Prior investigations have exploited flexible optical fibers to conduct light into and out of the deep brain, enabling the recording of neural dynamics while ensuring unrestricted animal behavior[12-14]. This fiber-based recording approach can be easily extended to multiple implantations for multi-site recording[15, 16], although the collected signals were averaged from many neurons in a volume. In addition, dense coherent optical fibers can be bundled for cellular resolution imaging[12, 13], and multi-region neuronal recording is possible when multiple fiber bundles are implanted[17]. However, these fiber-optic microscopes hold several inherent drawbacks that hinder their applications, such as limited spatial resolution, small field of view (FOV), low transfer efficiency of fluorescence, and unpleasant susceptibility to animal motion[18]. Moreover, a complicated well-designed commutator is needed to relieve torsional strain within the bundle as the mouse behaves[13].

Miniature head-mounted fluorescence microscopy is another class of imaging modality supporting cellular-resolution recording of neural activity in freely moving animals[19-21]. In contrast to the fiber-optics microscope, it integrates all its optical components within a small lightweight housing carried by animals. This design yields superior performance in terms of optical sensitivity, field of view, attainable resolution, mechanical flexibility, cost and portability[18]. The past decade has witnessed great success in exploring the cellular dynamics underlying behavior, cognition and sensation[20, 22, 23]. To enable multi-area recording, several works have customized designs to achieve large FOVs[24-27]. However, this strategy is mainly suitable for superficial cortical areas because implanting large GRIN lenses into subcortical areas may cause severe brain damage. Since the targeted regions are usually spatially distant and at different depths, simply increasing the FOV is not flexible enough to handle all experimental scenarios. Thus, a more practical solution is to image each brain region with a separate head-mounted microscope. However, the implantation of more than two microscopes on the mouse brain remains challenging due to constraints in available head space and the significant weight of the equipment [28, 29]. Recently, an optimized design reduced the weight from ∼1.9 g to 1.0 g[26], but the device size is still not ready for multiple-region recording on small animals such as mice and songbirds.

Here, we report the development of an open-source ultra-compact head-mounted microscope, named TINIscope, that has substantially reduced size and weight (0.43 g, Table 1), achieved through extreme optimization in optical, electrical and mechanical designs. As standalone equipment, TINIscope reaches a new level of miniaturization in head-mounted microscopes, reducing the burden added to smaller or developing animals like juvenile songbirds. Based on the TINIscope design, we further systematically developed an experimental paradigm, including multi-site implantations and a commutator for untangling, for multi-region calcium imaging in freely behaving mice. In proof-of-concept experiments, we achieved simultaneous neural activity recording of over 1000 neurons in four subregions of the hippocampus in free-moving mice and extracted neural activity with spatially modulated properties in these regions. A detailed analysis of recorded neural activity revealed clustered neuronal populations spanning all four subregions. The compact design of TINIscope enables flexible combination with optogenetic or electrophysiological tools to achieve a more versatile experimental design. In our experiments, we simultaneously collected individual neuronal activities in four hippocampal subregions in response to optogenetic or electrical stimulations in the anterior cingulate cortex (ACC). In addition, we jointly recorded calcium signals and local field potentials (LFPs) in each hippocampal subregion and analyzed population activity patterns concurrent with ripple onsets. These initial efforts suggest that TINIscope can be applied to recording neural activity in multiple brain regions at the single-cell level as well as integrating with a variety of techniques to become a multifunctional tool for exploring neural mechanisms.

**Table 1.**
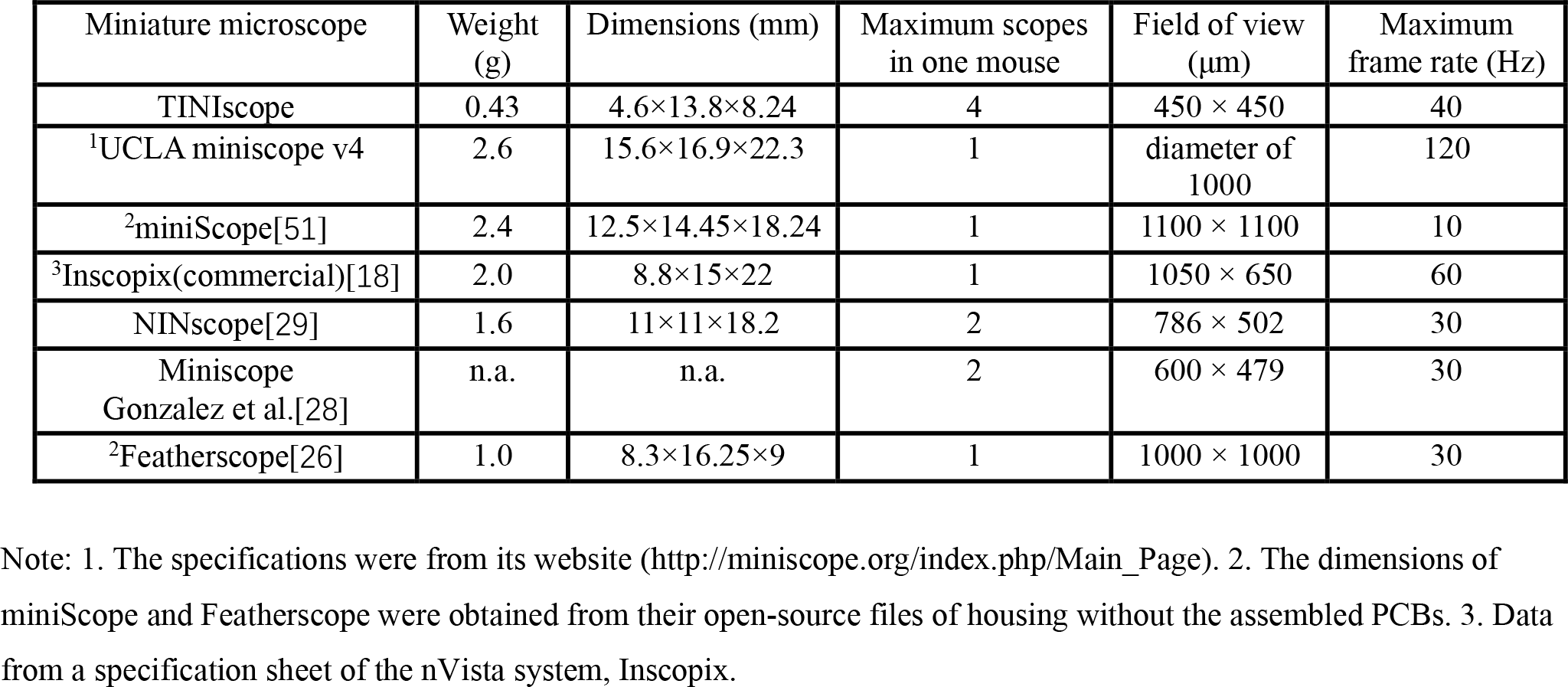
Key feature comparison of TINIscope and other miniature fluorescence microscopes

## RESULTS

### Design of the TINIscope system for multi-region recording

The key component of our multi-region recording system is the ultra-compact TINIscope mounted on the animal head. It uses a GRIN lens as its objective and a short-pass dichroic mirror to reflect emitted fluorescent light to an image sensor located on the side of the 3D-printed main body of the device (Fig. 1A, Supplementary Fig. 1). The fundamental optical pathway of TINIscope is the same as that of classical head-mounted microscopes[18, 20], whereas essential modifications were made to enable multi-region imaging with three design principles: compatibility with multiple implantations, compactness and lightness.

**Figure 1.**
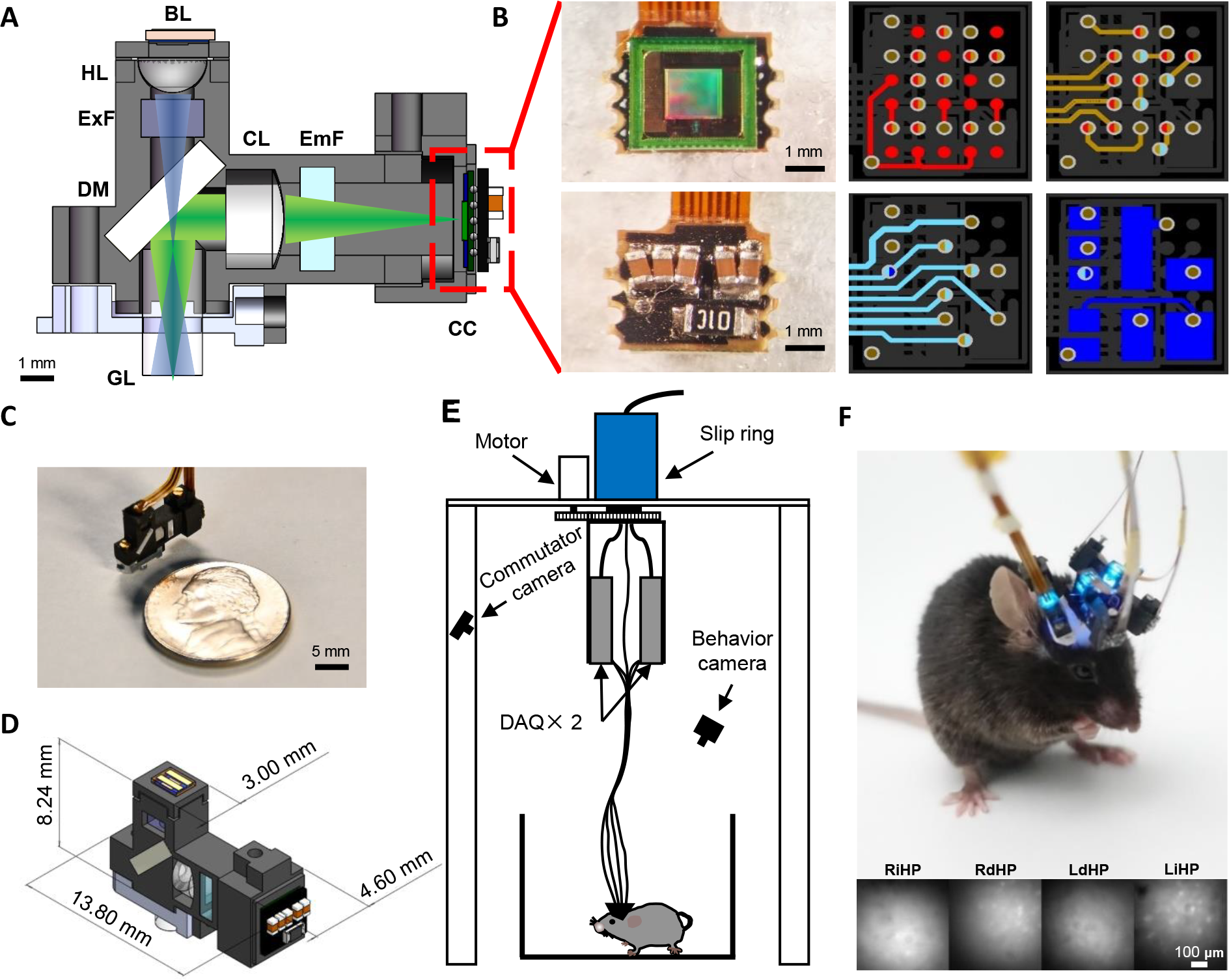
Tightly integrated neuronal imaging fluorescence microscope (TINIscope). **A**, Section diagram of TINIscope. BL, blue LED; HL, half-ball lens; ExF, excitation filter; DM, dichroic mirror; GL, GRIN objective lens; CL, convex lens; EmF, emission filter; CC, CMOS camera. **B**, Photos of an image sensor with HDI rigid PCB and layout of stacked layers (from top to bottom layer: red, brown, cyan and blue) with blind and buried vias (bicolor). **C**, Photo of TINIscope and a nickel shown for scale. **D**, dimensions of TINIscope. **E**, Schematic diagram of the experimental system with TINIscope. **F**, Top: photo of a mouse with 4 head-mounted TINIscopes. Bottom: Simultaneously recorded images of 4 hippocampal subregions.

First, the design of TINIscope deviates from the conventional setup, with the placement of the CMOS sensor on the side and LED excitation on the top (Fig. 1A, Supplementary Fig. 1). In the conventional setup, accommodating multiple pieces of equipment on the mouse head, especially closely located areas, is challenging due to the spatially conflicting large CMOS sensors in the upright direction. Since the LED excitation pathway is much smaller, flipping the excitation and emission pathways can alleviate this issue. Thus, we kept the illumination arm straight while using the dichroic mirror to reflect emission fluorescence to the side-placed CMOS imaging sensor. When 4 TINIscopes are implanted on the mouse brain, we can always rotate them to spatially distribute their CMOS imaging sensors. In practice, we designed a virtual simulation pipeline for arranging equipment to guide our surgical implantation (Supplementary Note 1).

Second, TINIscope incorporates an ultra-compact light pathway to match the selected smaller LED device (1.3 mm × 1.7 mm) and CMOS imaging sensor (pixel size of 3 μm). In the illumination arm, a half-ball lens (HL) with a diameter of 2 mm is used to collect excitation light from the LED. This design not only matches the small LED size but also allows for a shorter distance between the HL and the GRIN lens (GL), resulting in a further reduction in equipment size. Zemax optical modeling showed that an HL-GL distance of 6 mm can illuminate a 0.63 mm-diameter area (Supplementary Fig. 1A). Additionally, TINIscope employs CMOS sensors with a small pixel size (3 μm), allowing a shortened pathway with a smaller magnification factor (∼2.4) that can achieve comparable spatial resolution and reasonably smaller FOV (∼450 μm × 450 μm) compared to those of existing systems (Supplementary Fig. 1B-G).

Third, the image sensor directly transmits serialized data without utilizing a serializer chip, which was a major component in previous head-mounted scopes, thereby simplifying the peripheral circuit of the PCB (Fig. 1B, Supplementary Fig. 2). Furthermore, other components, such as the power supply chip and oscillator, were removed from the circuit, leaving only essential capacitors and one resistor on the board (Fig. 1B right and Supplementary Fig. 2A, B). A customized HDI rigid-flex PCB was subsequently developed to further reduce the size of the head-mounted PCB and enable flexible connectivity to the DAQ board (Supplementary Note 2). Following this, mechanical modifications were implemented to enhance the compatibility of the head-mounted microscope with multi-site implantations (Fig. 1 C, D and Supplementary Note 3).

In TINIscope, multiple image sensors are connected to the data acquisition (DAQ) board through flexible PCB cables. To mitigate the possibility of entanglement caused by the motion of the mouse, a commutator was developed (Fig. 1E). The commutator employs a stepper motor to rotate an electrical slip ring and unravel any twisted cables according to their shapes monitored by a camera (Supplementary Fig. 4).

With such a configuration, the head-mounted module of TINIscope weighs only 0.43 g, which is substantially smaller and lighter than all existing devices while preserving image quality (Fig. 1F, Table 1 and Supplementary Video 1). Meanwhile, our quantitative comparisons substantiated that the installation of multiple TINIscopes had no notable impact on the mobility of mice (Supplementary Figure 5). Furthermore, a multi-session studies (days 1, 3, and 5) verified the feasibility of imaging and registering the same population of neurons with TINIscope (Supplementary Figure 6). This compact design makes it possible to implant up to 4 devices into a mouse brain (Fig. 1F, Supplementary Fig. 3D).

### Recording of four hippocampal subregions with TINIscope

The hippocampus is an essential brain area involved in spatial navigation, learning and memory, and mental disorders, and subregions of the hippocampus have been shown to be functionally and anatomically distinct. We therefore conducted concept experiments to investigate hippocampal neural coding in freely behaving mice with TINIscope, aiming to validate the feasibility of simultaneous four-region recording in a mouse (Fig. 2A). Four subregions of the bilateral hippocampi, including the right intermediate hippocampus (RiHP), right dorsal hippocampus (RdHP), left dorsal hippocampus (LdHP), and left intermediate hippocampus (LiHP), of 8-week-old mice were selected for infection with an adeno-associated virus (AAV2/9-hSyn-GCaMP6s) to express the calcium-sensitive fluorescent protein GCaMP6s. Three weeks later, a GRIN lens was implanted in each of these regions. Viral infection and lens implantation were later verified through examination of brain slices (Fig. 2B). The subsequent experiments presented in the main text employed the same recording sites and basic experimental procedures.

**Figure 2.**
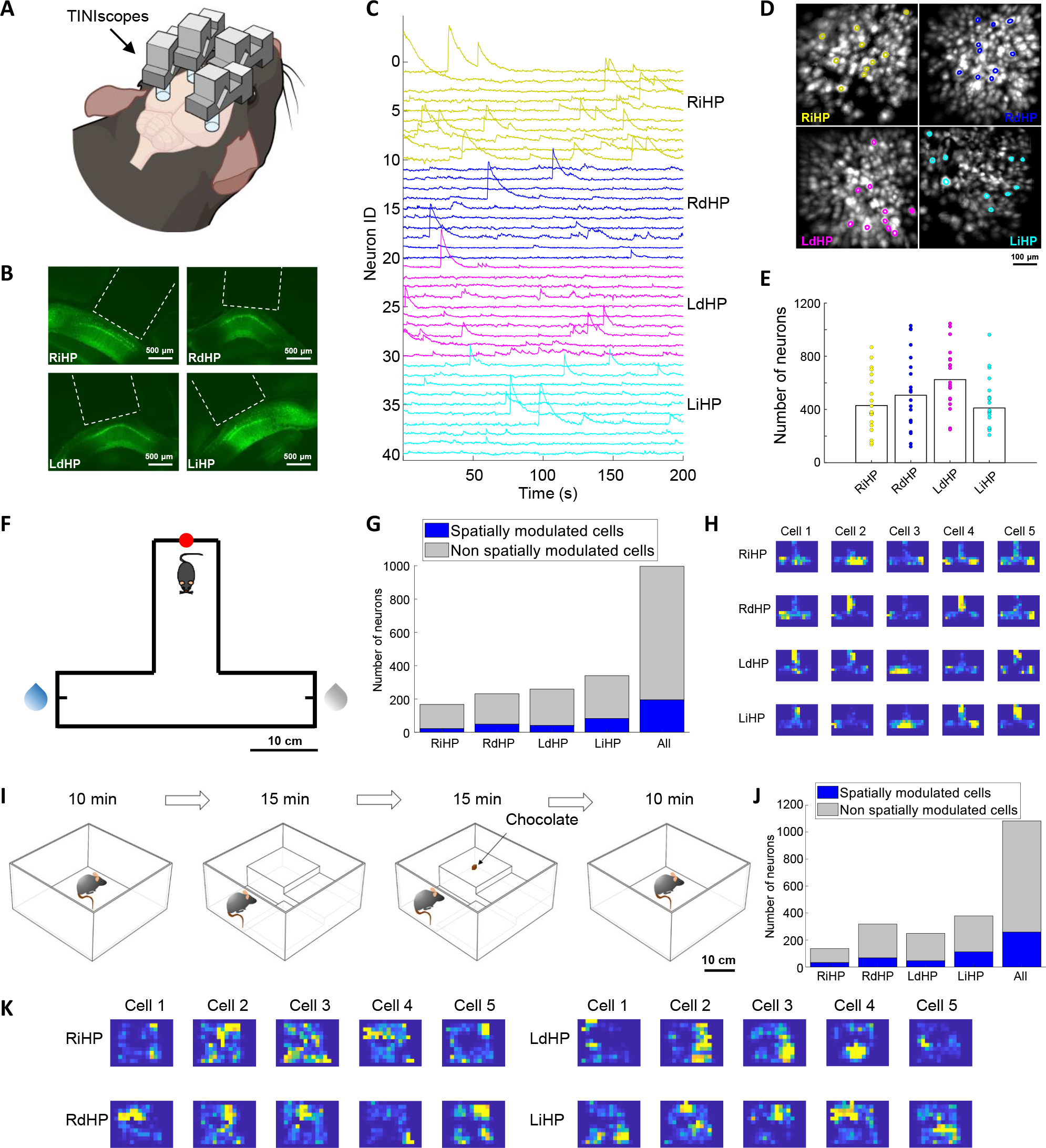
Simultaneous calcium imaging of 4 hippocampal subregions in different behavioral experiments. **A**, Diagram of 4-region recordings in mice. **B**, Fluorescence images of brain slices around the four recorded hippocampal subregions. The white dashed lines indicate GRIN lenses. **C**, Calcium traces of example neurons from the RiHP (yellow), RdHP (blue), LdHP (magenta), and LiHP (cyan). **D**, The spatial contours of example neurons in C and the corresponding maximum intensity map (MIP) of the background-subtracted videos. **E**, Number of identified neurons in all sessions and their averages (*n* = 20 sessions, 6 mice). **F**, Paradigm of the T-maze task. The water reward was given at a random end in each trial. Red dot: start point; blue droplet: reward side; gray droplet: non-reward side. **G**, The proportion of spatially modulated cells during the T-maze task. **H**, Place fields of example spatially modulated cells in the T-maze box. **I-K**, Same as **F-H**, but the mouse was in an open field exploration experiment. The mouse explored freely in an open field with changing environments.

After two weeks of recovery and adaptation, simultaneous cellular-resolution imaging of neural activities was conducted using 4 separate TINIscopes while the mouse was moving freely in various behavioral paradigms (Fig. 2F, I and Supplementary Video 2). Subsequently, the CNMF-e[30] method was employed to identify neurons and extract their temporal traces from the raw videos (Fig. 2C, D and Supplementary Video 2). The field of view (FOV) of each region was ∼450 μm × 450 μm and contained a few hundred neurons (Fig. 2E, *n* = 20 sessions from 6 mice; the minimum numbers of recorded neurons in RiHP, RdHP, LdHP, and LiHP were 137, 121, 249, and 208 respectively, while the maximum numbers were 868, 1030, 1046, and 962 respectively).

To elucidate the spatial encoding properties of the hippocampus, experiments were conducted in a T-maze and a modified open field arena. During the T-maze test, a water-deprived mouse was required to enter the starting zone located at the end of the vertical arm to initiate a trial. Upon entering the starting zone, an LED would be illuminated, and one of the two ends of the horizontal arms would be randomly designated as the available option for water. The mouse was then allowed to freely explore the T-maze until they obtained a reward, which signified the completion of a trial (Fig. 2F). We employed mutual information analysis to identify neurons exhibiting spatial tuning. All recorded regions contained spatially tuned neurons with a proportion of approximately 1/5. Fig. 2F-H shows a mouse with 22/167, 49/231, 41/259 and 83/340 spatially tuned neurons in the RiHP, RdHP, LdHP, and LiHP, respectively.

During the modified open field test, we introduced steps of varying heights and a small piece of chocolate into the standard open field arena, providing mouse with a dynamic and changing environment for exploration (Fig. 2I). The mouse, equipped with four recording devices, was able to freely navigate and explore different contexts (Supplementary Video 2), demonstrating that the lightweight TINIscope had no obvious impacts on their behavior. This further supports the suitability of TINIscope for recording neural activity across multiple brain regions. Fig. 2I-K shows the same mouse with 33/137, 68/318, 46/249, and 112/379 spatially tuned neurons in an open-field environment in stages 2 and 3. These results indicate that the hippocampus exhibits widespread encoding of spatial information across subregions.

### Multi-modal experiment design with TINIscope

To understand how brain-wide neural activity supports complex behaviors, it is usually necessary to employ multiple complementary modalities for recording or manipulating neuronal activity during behavior. The compact dimensions and lightweight construction of TINIscope facilitate effortless combination of optical or electrophysiological modules, without incurring significant weight or size constraints (Fig. 3A, Supplementary Videos 3 and 4). To demonstrate this advantage of TINIscope, we first implanted two optical fiber cannulas and four recording devices in the bilateral ACC with ChrimsonR expression (Supplementary Fig. 7A) and four hippocampal subregions with GCaMP6s expression, respectively, to investigate their induced neuronal activity in the hippocampus (Fig. 3B). We observed an increase in the mean fluorescence of each region, as well as the activity of certain individual neurons following stimulation (Fig. 3C, D and Supplementary Fig. 7), in agreement with the reported ACC-hippocampal connections[31, 32].

**Figure 3.**
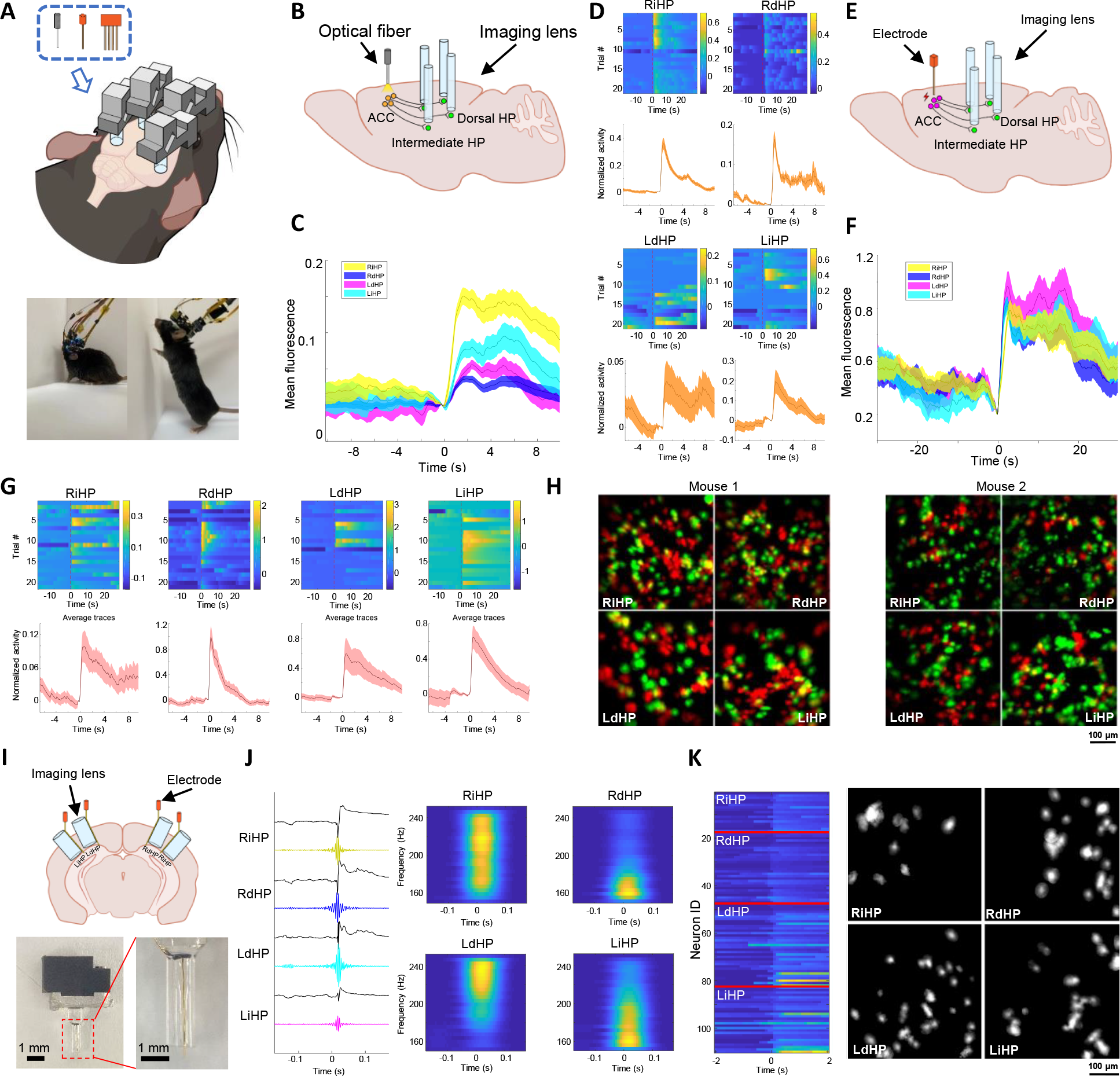
Combination of optogenetic/electrophysiological modules and TINIscope. **A**, Top: diagram of potential experimental paradigms that combine multiple-TINIscope imaging and other technique modules, as listed in the dashed box (optogenetics, electrical stimulation and electrophysiological recordings). Bottom: photos of mice carrying 4 TINIscopes together with 2 electrical stimulating electrodes (left) or 4 extracellular recording electrodes (right). **B**, Diagram of investigating the ACC-HP circuit by combining optogenetics with 4-region TINIscope imaging. **C**, Mean of the background-subtracted fluorescence signals averaged over all left-ACC stimulation trials (*n* = 21 trials, 1 mouse, t=0 indicates the stimulation onset). **D**, Normalized activity of example ACC-responsive neurons in different trials (top) and the corresponding mean activity over trials (bottom). The traces were normalized using the estimated noise level and centered around the value at the onset of stimulation (t=0). **E-G**, Same as **B-D**, but replacing optogenetic stimulation with electrical stimulation (*n* = 42 trials, 2 mice). **H**, Left: spatial footprints of neurons responding to left (red) or right (green) ACC stimulation, respectively; right: same as the left but with the other mouse. I, Top: illustration of joint calcium imaging and LFP recording in 4 hippocampal subregions. Bottom: Photo of the electrode-lens complex. **J**, Raw LFP and filtered signals (150-250 Hz) when all 4 regions show SWRs together. Right, spectrograms of LFP signals in 4 regions. **K**, Synchronous calcium traces (left) and their spatial footprints (right) concurrent with the SWR in **J**. Shaded areas in **C, D, F, J** correspond to the mean±s.e.m.

To further validate the neural activity induced by this optogenetic stimulation and to explore the possibility of integrating TINIscope with the electrical stimulation module, we bilaterally implanted electrodes in the ACC and recording devices in four subregions of the hippocampus (Fig. 3E-G, Supplementary Fig. 8). Each stimulation cycle started with 1 second of 30 Hz 300 μA current stimulation and then rested for 60 seconds. The mean fluorescence of all regions significantly improved after stimulation (Fig. 3G and Supplementary Fig. 8D, E, *P* < 0.001, Wilcoxon matched-pairs sign rank test). After extracting all neuronal activity, we identified stimulation-responsive neurons by comparing their mean activities 2 seconds before and after stimulation (Fig. 3G). We repeated these experiments on two mice and observed that all imaged regions contained a proportion of neurons that responded to stimulations on either side of the ACC (Fig. 3H). These results suggest that TINIscope is ideal for seamless combination with established brain manipulation methods, enabling comprehensive investigation of functional connectivity within neural circuits.

Due to the slower response of neuronal calcium signals compared to electrical signals, the high-frequency component of neural activity is lost in calcium imaging recording. To overcome these limitations, a viable approach is to simultaneously perform calcium imaging and electrode recording at the same site. To investigate the feasibility of using TINIscope for calcium imaging in conjunction with electrode recordings in the same brain region, we co-implanted the GRIN lens and electrodes in four hippocampal subregions (Fig. 3I). TINIscope provided calcium signals with single-cell resolution, while the electrodes enabled the acquisition of electrical signals with high temporal resolution. With this configuration, we successfully conducted electrophysiological recordings concurrently with TINIscope imaging at all four recording sites (Supplementary Fig. 9A, B). By filtering the LFPs, we were able to reliably detect signal patterns such as sharp-wave ripples (SWRs), a distinct waveform that propagates in the hippocampus[33, 34], in all recorded regions (Fig. 3J, Supplementary Fig. 9C, D). In conjunction with the results from TINIscopes, we were able to investigate the firing patterns of neurons within multiple regions, coinciding with SWRs (Fig. 3K, Supplementary Fig. 9C, E). The spatial organization of these synchronized neurons may yield valuable insights into the formation and propagation of SWRs.

### Decoding mouse position from extracted neuronal activity

In our previous experiments, we observed a substantial amount of spatially modulated neurons in the recorded hippocampal subregions. Thus, we further explored how spatial information is distributed in those regions. To address this inquiry, we trained a machine learning model to decode mouse locations from neuronal activities[35]. The LSTM algorithm with specific hyperparameters (units = 200, dropout = 0.25, number of training epochs = 10) was selected because it has empirically higher decoding accuracy for hippocampus data. Prior to decoding, the extracted calcium traces were temporally aligned with mouse locations, and silent periods at the start or end of trials were manually removed. The rest of the data were equally split into 10 folds, with 9 folds used for training an LSTM decoder and the last fold used for calculating the prediction error using a 10-fold cross-validation procedure (Fig. 4A).

**Figure 4.**
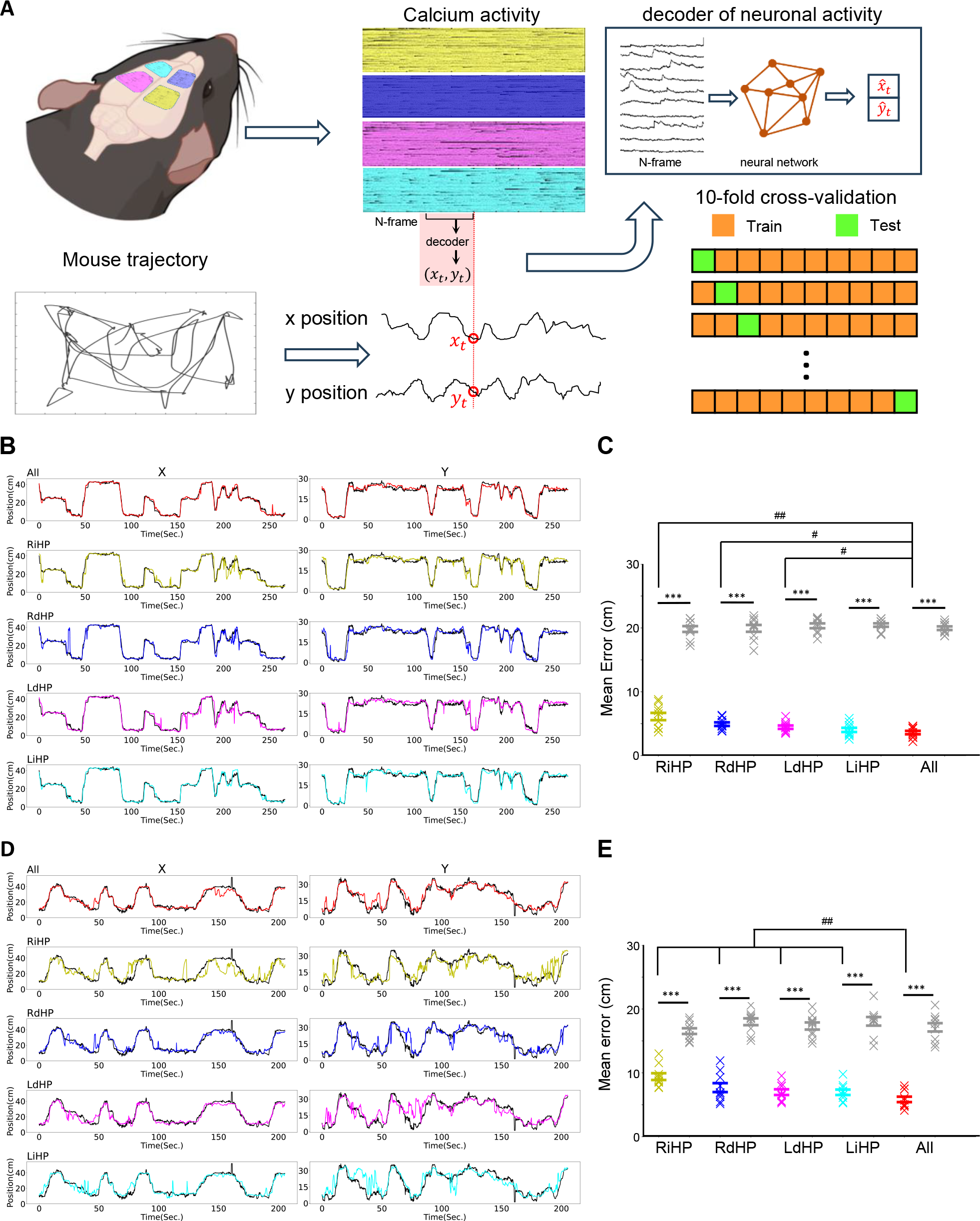
Decoding mouse position from extracted neuronal traces. **A**, Schematic illustration of machine-learning-based decoders for predicting mouse position from neuronal activity. Bottom right: the extracted calcium traces were split into 10 folds, and one testing fold was sequentially chosen to calculate the decoding error with the decoder trained from the remaining 9 folds. Decoding the position at each time point requires the population neuronal activity of the previous N (here, N = 5) frames. The LSTM network was used as the decoding model in this work. **B**, Example decoded mouse positions (color lines) on testing data and the true position (black lines) in the T-maze experiment. The number of temporal bins in this testing fold is 2670 (bin size=100 ms). **C**, The mean decoding error using different sets of neuronal traces in the T-maze experiment. Gray data points correspond to the chance level decoding errors where the mouse positions were randomly shuffled. \*\*\**P* < 0.001, Mann-Whitney test, *n* = 10 folds, 1 mouse. ##*P* < 0.01, #*P* < 0.05, Wilcoxon matched-pairs sign rank test, *n* = 10 folds, 1 mouse. The results show the mean ± s.e.m. **D** and **E**, same as **B** and **C**, but the mouse was in the open field experiment (temporal bins = 2059).

By examining the decoding results from a representative fold, we observed that across both T-maze and open field paradigms, the integration of neuronal activities across multiple brain regions resulted in superior prediction of mouse locations compared to utilizing activities from a single brain region alone (Fig. 4B, D and Supplementary Video 5). To determine the chance level of decoding error, the same procedure was performed on location-shuffled data, where the location vector was flipped in time and randomly cycle-shifted for at least 2000 frames. All decoding results exhibited significantly better performances above the chance level (Fig. 4C, E).

During the T-maze exploration, we observed that neuronal activities from a single brain region accurately decoded the mouse’s position, with a median prediction error approximately equal to the length of the mouse body. However, during a more random and complex exploration within the open field, relying solely on neuronal activities from a single brain region was no longer sufficient to accurately decode the mouse’s position. Instead, the participation of neuronal activities from multiple recorded regions significantly improved the decoding accuracy, enabling a more precise estimation of the mouse’s position (*n* =10 folds in one mouse, cross-validation, Wilcoxon’s rank-sum test, Fig. 4C, E, with median errors of 6.13 ± 0.57 cm in RiHP, 4.94 ± 0.28 cm in RdHP, 4.46 ± 0.26 cm in LdHP, 4.03 ± 0.32 cm in LiHP, and 3.62 ± 0.26 cm in all data in the T-maze and 9.41 ± 0.53 cm in RiHP, 7.69 ± 0.70 cm in RdHP, 6.98 ± 0.45 cm in LdHP, 6.95 ± 0.42 cm in LiHP, and 5.84 ± 0.43 cm in all data in the open field). Overall, these results suggest that the encoded spatial information is distributed in a larger area than the FOV range of a single scope, consistent with previous studies[36].

### Multiple-region recording reveals distributed activity patterns

Dynamic activation of neuronal assemblies is considered a key mechanism underlying cognition and behavior[34, 37, 38]. TINIscope provides a great opportunity for studying behavior-relevant assemblies that span multiple brain regions. Hence, we proceeded to investigate the formation of neuronal assemblies among the recorded neurons and examine their correlation with the spatial information of the mouse.

In the T-maze and open field tests, we analyzed potential assembly dynamics by identifying synchronous calcium events (SCEs), clustering them, and assigning single neurons to putative assemblies (Fig. 5A-D and Supplementary Fig. 10A-C). Assemblies were identified using the procedure described in an early hippocampus study[34]. This method first detected SCEs where at least M neurons were active (value of z-scored calcium trace > 1) for more than 500 ms. M was determined by traversing a wide range of candidate values and selecting the one that maximized the number of detected SCEs (note: a small value of M results in longer but fewer SCEs). Once detected, each SCE used a binary vector to indicate the status of all neurons within that SCE. Then, a two-stage robust K-means clustering algorithm (Euclidean distance) was applied to cluster these SCE vectors. In stage 1, 100 conventional K-means were applied to find K cluster centers, while in stage 2, these 100 cluster centers were clustered again using K-means clustering, yielding stable cluster centers. The value of K was chosen between 2 and 10, and the one with the best average silhouette value was used. Afterwards, each neuron was defined as participating in an SCE cluster if the neuron showed a significantly higher activation rate (above the 95th percentile of reshuffled data) in the SCEs of that cluster. In this way, each SCE cluster was associated with a cell assembly comprised of neurons participating in that cluster.

**Figure 5.**
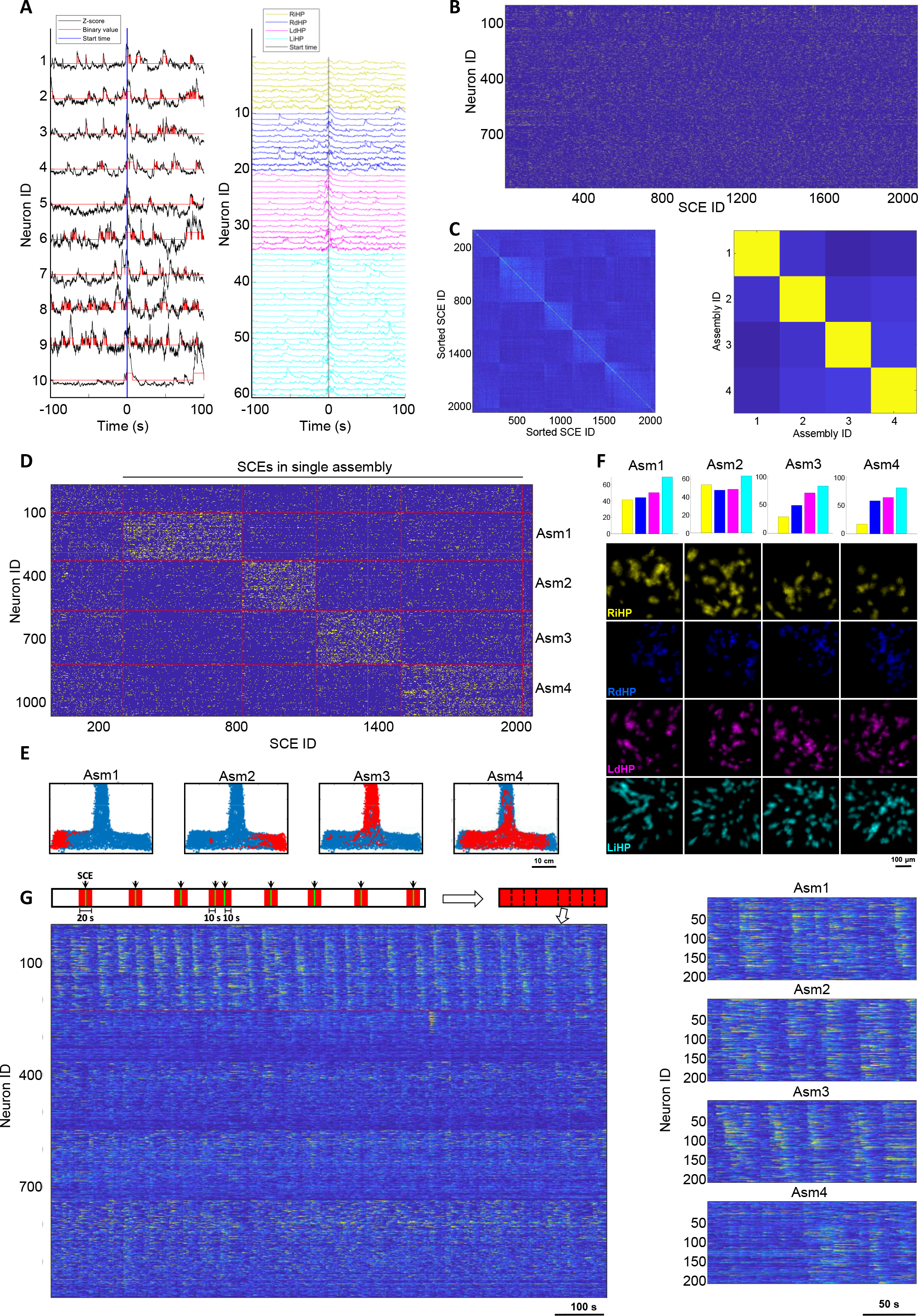
Neuronal assemblies during the T-maze task. **A**, Detection of SCEs from extracted calcium traces. Left: activities of the example neurons were z-scored and thresholded at 1 standard deviation (SD); right, neuronal activity of all neurons participating in an SCE. The vertical lines indicate the onset of an SCE. **B**, Raster plot of neuronal participation in all SCEs. **C**, Correlation maps of the detected SCEs (left) and the identified neuronal assemblies (right). The SCE IDs were ordered by the associated assemblies of the participating neurons in each SCE. **D**, Same as B but the neuron IDs and SCE IDs were ordered to match the found assemblies. The SCEs to the left column were not involved in any assembly, while the SCEs to the right column were involved in multiple assemblies. **E**, The mouse’s position when SCEs associated a specific assembly occurred. **F**, The number of neurons in each region (color coded as labels below) and their spatial footprints in different assemblies. **G**, Neurons associated with each assembly showed repeated activation sequences. Left is the concatenated calcium traces, as illustrated in the top diagram. Only time intervals within ±10 second windows of all SCEs associated with an example assembly (assembly 3, including neurons above the red line) will be preserved. Neurons were grouped by their associations with different assemblies. Within each assembly, neuron IDs were sorted according to their peak intensity time. Right, repeated activation sequences of neurons associated with each assembly.

SCEs were grouped afterwards based on the assemblies they activated. We proceeded to generate plots of mouse locations corresponding to the occurrences of these neuronal assemblies, without any prior knowledge of their actual locations. In the T-maze task, assemblies 1-3 exhibited a clear preference for specific locations, whereas assembly 4 did not show such a preference (Fig. 5E). Notably, these neuronal assemblies were spatially distributed throughout all the recorded brain regions (Fig. 5F). The assemblies associated with explicit location preference displayed recurring firing sequences, while the other assemblies did not exhibit such recurring patterns (Fig. 5G). Similar results were obtained in the open field experiment (Supplementary Fig. 10D-F), suggesting a brain-wide dynamic population encoding of environmental information.

## DISCUSSION

The neuroscience community has an urgent demand for tools that can perform concurrent cellular-resolution calcium imaging in multiple brain regions of freely behaving animals. In this work, we have demonstrated the applicability of TINIscope to meet these requirements, enabling fundamentally new explorations of the brain-spanning coordinated cellular dynamics that underlie sensation, cognition, and action. TINIscope represents the lightest and smallest head-mounted miniature microscope with a large margin. In the context of single region recordings, employment of TINIscope will yield a remarkable decrease in the burden placed on very small and developing animals when compared to other microscopes within its class.

To achieve the goal of multi-region recording, a comprehensive system design is necessary, which involves more than simply reducing the size and weight of the microscope. In this work, considerable efforts have been directed toward optimizing the optical path, housing, and baseplates to enable the spatial arrangement of multiple TINIscopes. Additionally, a virtual simulation workflow was developed to guide our implantation surgery. Our demonstration experiments successfully recorded four closely situated hippocampal subregions with severe spatial limiting issues, thereby indicating support for nearly any combination of four brain regions in mice. To avoid potential entanglements of flexible PCBs, we also developed a customized commutator for our experimental system. Moreover, we developed our data acquisition system and GUI software to streamline data collection. The entire system’s design is open source and can be further customized to include additional functionalities such as e-focus, dual-color imaging, in situ optogenetics stimulation, and volumetric imaging[39]. The technical approaches proposed for existing miniature microscopes, such as μTlens for z-focusing[40], dual excitation light sources[41], interleaved readout of CMOS sensors, and micro lens array for volumetric imaging[42], can be adapted for TINIscope. The primary challenges lie in integrating these techniques into the ultra-compact design of TINIscope and customizing the necessary electronic components to support them.

The investigation of numerous neuroscience questions necessitates the use of multimodal combinations of recording and manipulating tools. Our TINIscope provides a much-needed toolkit for addressing neuroscience questions regarding multi-region interactions during specific brain functions, with its ultra-compact design facilitating integration with other tools. We validated its easy combination with classic tools for optogenetics, electrical stimulation and LFP recordings, providing insights into factors related to imaged calcium activity. The family of head-mounted fluorescence microscopes is rapidly expanding[19], with TINIscope being a new member featuring the smallest size and lightest weight. We expect to see future experimental designs combining TINIscope with other members noted for large FOV[24-27]or multi-photon imaging[43-46].

Given the rapid advances in our anatomical knowledge of brain-wide neuronal connectivity[47, 48], TINIscope is a timely and valuable tool for enhancing our understanding of functional neuronal connectivity during complex behaviors and cognitive tasks performed in unrestrained conditions. We expect that our technique will help pave the road toward understanding brain circuit function from a more holistic perspective in conjunction with novel computational methods to extract insights from this wealth of information [2, 49, 50].

## MATERIALS AND METHODS

For detailed materials and methods, please see the supplementary data.

## ACKNOWLEDGEMENTS

We thank X. Tang and D.X. Wang for drawing diagrams in this paper and many helpful discussions for analyzing the data. We especially thank S.M. Zhang and L.F. Zhang for discussing the surgical procedure for calcium imaging. We thank C. Xu and C. Wang for discussing the interpretation of imaged data in the hippocampus, X.X. Wei and X.Y. Deng for insightful discussion, Jiangsu Brain Medical Technology Co. for supplying the MINIscope system for comparison of preliminary experimental results during the early design period of TINIscope, Y.F. Zhou and Y.F. Wang for helping to test the electronic system of TINIscope, Y. Shen, Q.Q. Liu for imaging the brain slices of experimental mice and C.H. Jia for preliminary test of TINIscope using calcium fluorescence dye.

## FUNDING

This work was supported by the Strategic Priority Research Program of the Chinese Academy of Sciences (XDB32030200), NSFC-GuangdongJointFund-U20A6005, NSFC (Grant No. 32100903), and the Key-Area Research and Development Program of Guangdong Province (2018B030331001).

### Conflict of interest statement

The University of Science and Technology of China has filed a patent application related to the design of TINIscope, for which F.Xue, K.Z., F.L., Q.Z., P.-M.L. and G.-Q.B. are named inventors. The remaining authors declare no competing interests.

## AUTHOR CONTRIBUTIONS

F.Xue, Q. Z., P.-M.L., G.-Q.B. conceptualized the project. F.Xue led the project under the supervision of P.Z., M.S. and G.-Q.B. F.Xue, K.Z. and Y.W. designed the TINIscope system. K.Z., F.L., P.Z. and D.Z. designed the experimental paradigm. F.L. and K.Z. conducted the surgery of mice, lens implantation and collection of image data. L.D. and X.Z. developed the GUI. F.Xue, P.Z. and K.Z. processed and analyzed the data. P.Z., F.Xu, Q.Z., D.Z., M.S., P.-M.L. and G.-Q.B. provided valuable insights for this project. F.Xue, P. Z., F.Xu, P.-M.L. and G.-Q.B. wrote the manuscript with input from all authors.

