## Supplementary Information for "Multi-region calcium imaging in freely behaving mice with ultra-compact head-mounted fluorescence microscopes"

`

*These authors contributed equally

This PDF file includes:

- Supplementary Figure 1-10
- Supplementary Note 1-3
- Materials and Methods
- Captions for Supplementary Video 1-6

### Content

SUPPLEMENTARY FIGURES ... 3
- Supplementary Figure 1. Optical design of TINIscope. ... 3
- Supplementary Figure 2. Electronic design of TINIscope. ... 5
- Supplementary Figure 3. Design of TINIscope arrangement in multiple-region implantations. ... 7
- Supplementary Figure 4. Design of the commutator system. ... 8
- Supplementary Figure 5. Comparison of moving ability of mice carrying no device, 4 TINIscopes, or 4 TINIscope & optical fiber. ... 9
- Supplementary Figure 6. Longitudinal calcium imaging with TINIscope in hippocampus. ... 10
- Supplementary Figure 7. Simultaneous imaging of 4 hippocampal subregions in response to optogenetic ACC stimulation. ... 11
- Supplementary Figure 8. Simultaneous imaging of 4 hippocampal subregions in response to electrical ACC stimulation. ... 13
- Supplementary Figure 9. Simultaneous LFP recording and calcium imaging in multiple brain regions. ... 14
- Supplementary Figure 10. Neuronal assemblies during open-field exploration. ... 15

SUPPLEMENTARY NOTES ... 16
- Supplementary Note 1 Placement of multiple TINIscopes via computer simulation ... 16
- Supplementary Note 2 Design of HDI rigid-flex PCB of TINIscope ... 16
- Supplementary Note 3 Mechanical design of TINIscope ... 16

MATERIALS AND METHODS ... 17

CAPTIONS FOR SUPPLEMENTARY VIDEO ... 22

### SUPPLEMENTARY FIGURES

**Supplementary Figure 1. Optical design of TINIscope.**

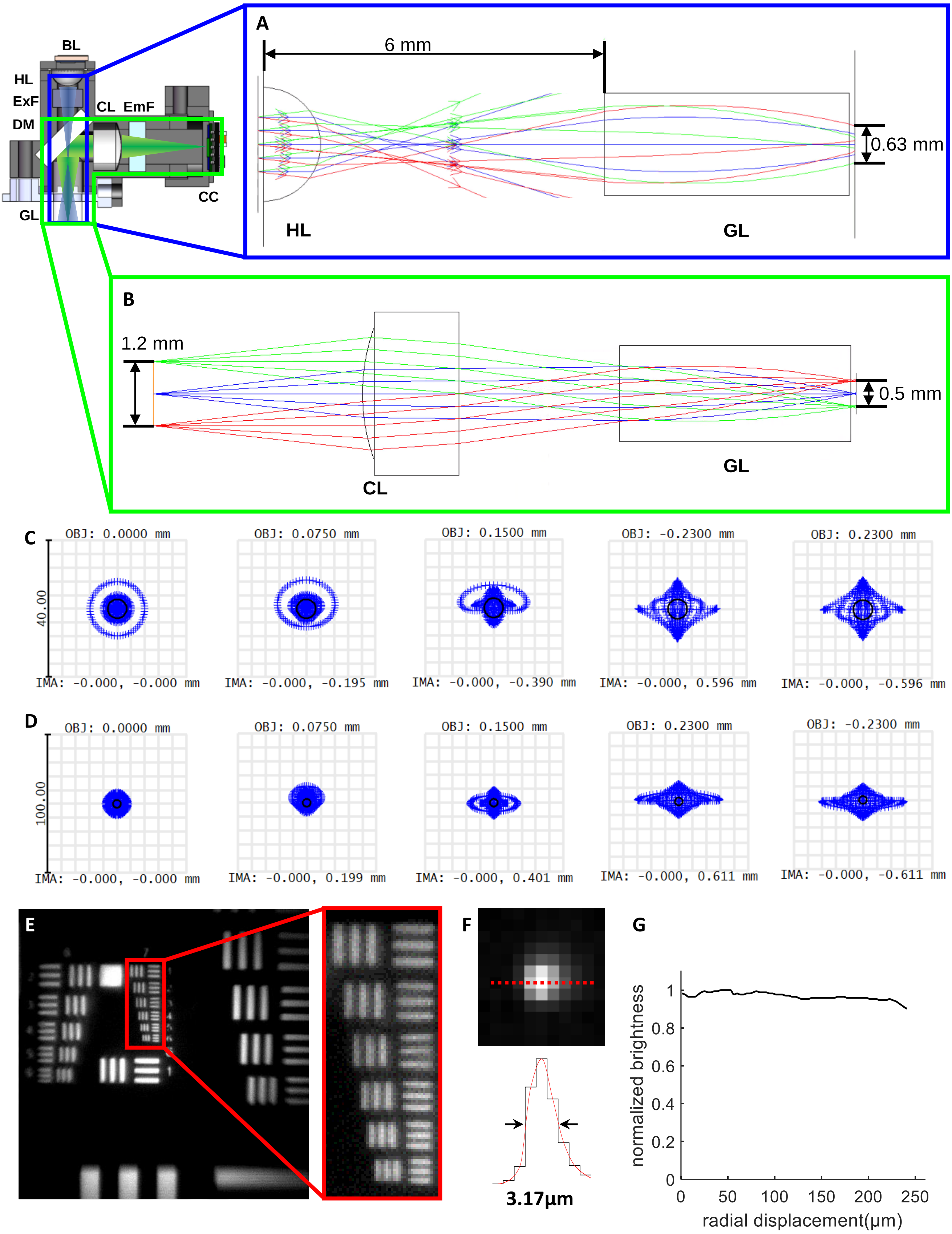

**Supplementary Figure 1. Optical design of TINIscope. A**, Illustration of the excitation light path in the illumination arm, demonstrating that the compact design (6 mm distance between HL and GL) allows for a large effective illumination area (diameter = 0.63 mm). BL, blue LED; HL, half-ball lens; ExF, excitation filter; DM, dichroic mirror; GL, GRIN objective lens; CL, convex lens; EmF, emission filter; CC, CMOS camera. **B**, Illustration of the emission light path in the imaging arm, with a magnification factor of 2.4 applied to the field of view (FOV) on the CMOS image sensor. **C**, Spot diagram of the TINIscope imaging system. **D**, Spot diagram of the TINIscope when utilizing a 1 mm relay lens. **E**, Image taken with the TINIscope of a USAF 1951 resolution target. The smallest line width in the red panel is 2.19 μm. **F**, Top: image of a 1 μm diameter fluorescence bead captured with TINIscope system. Bottom: line profile along the x axis of the bead image and the full width at half maximum (FWHM). **G**, Radial profile of the illumination at the focal plane.

**Supplementary Figure 2. Electronic design of TINIscope.**

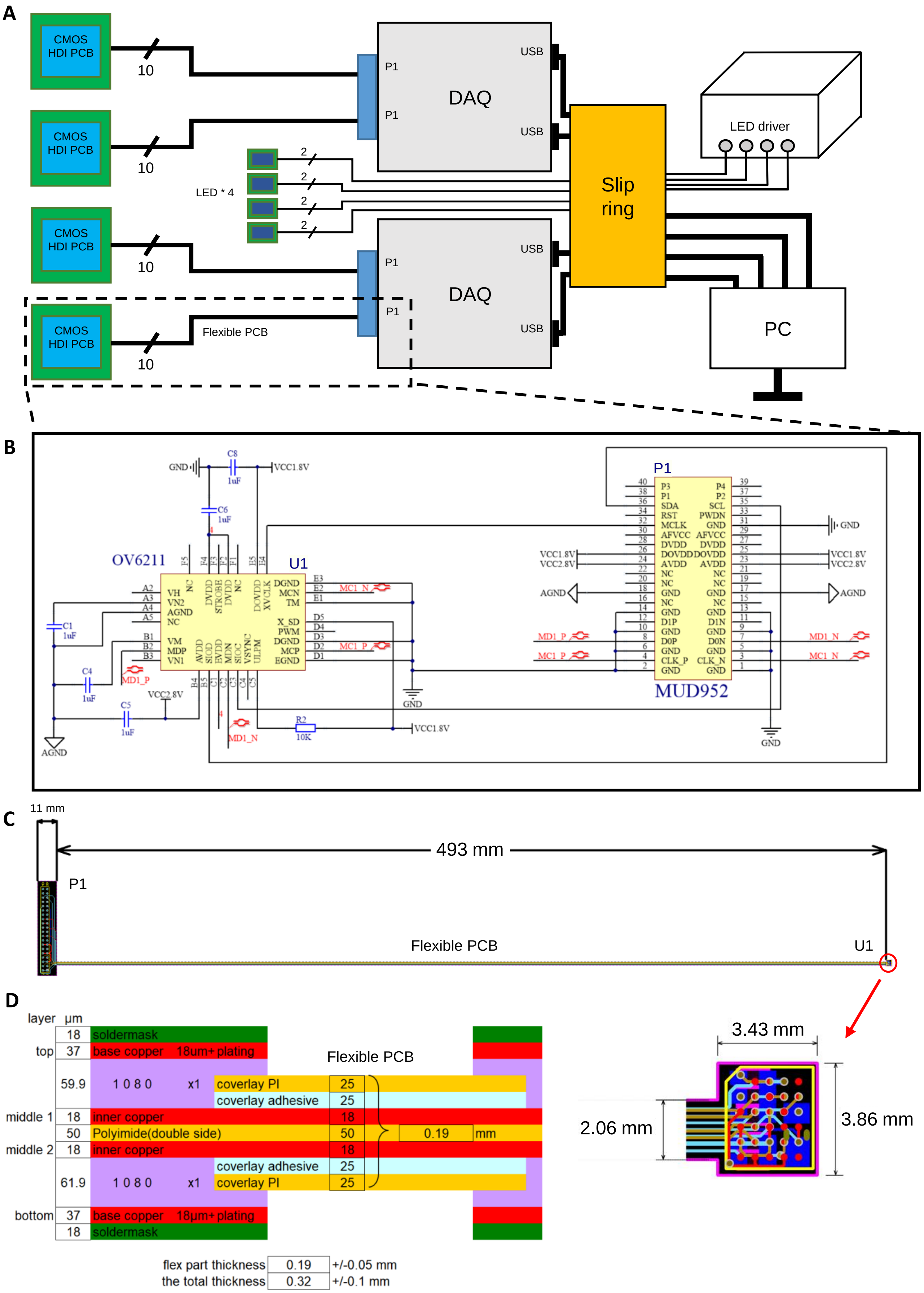

**Supplementary Figure 2. Electronic design of TINIscope. A**, Diagram of the electronic system for data acquisition and LED control. The CMOS image sensor signals are transmitted to the data acquisition (DAQ) boards via a flexible PCB cable and then transmitted to a computer. Each LED light is controlled by a driver with a 2-wire connection. **B**, Electronic schematic between the high-density interconnect (HDI) PCB and DAQ board in TINIscope. The flexible PCB cable is omitted for clarity. **C**, Layout and dimensions of the rigid-flex HDI PCB. The arrow indicates the magnified layout of the HDI image sensor PCB in the red circle. **D**, Layer stack indicating PCB connections.

**Supplementary Figure 3. Design of TINIscope arrangement in multiple-region implantations.**

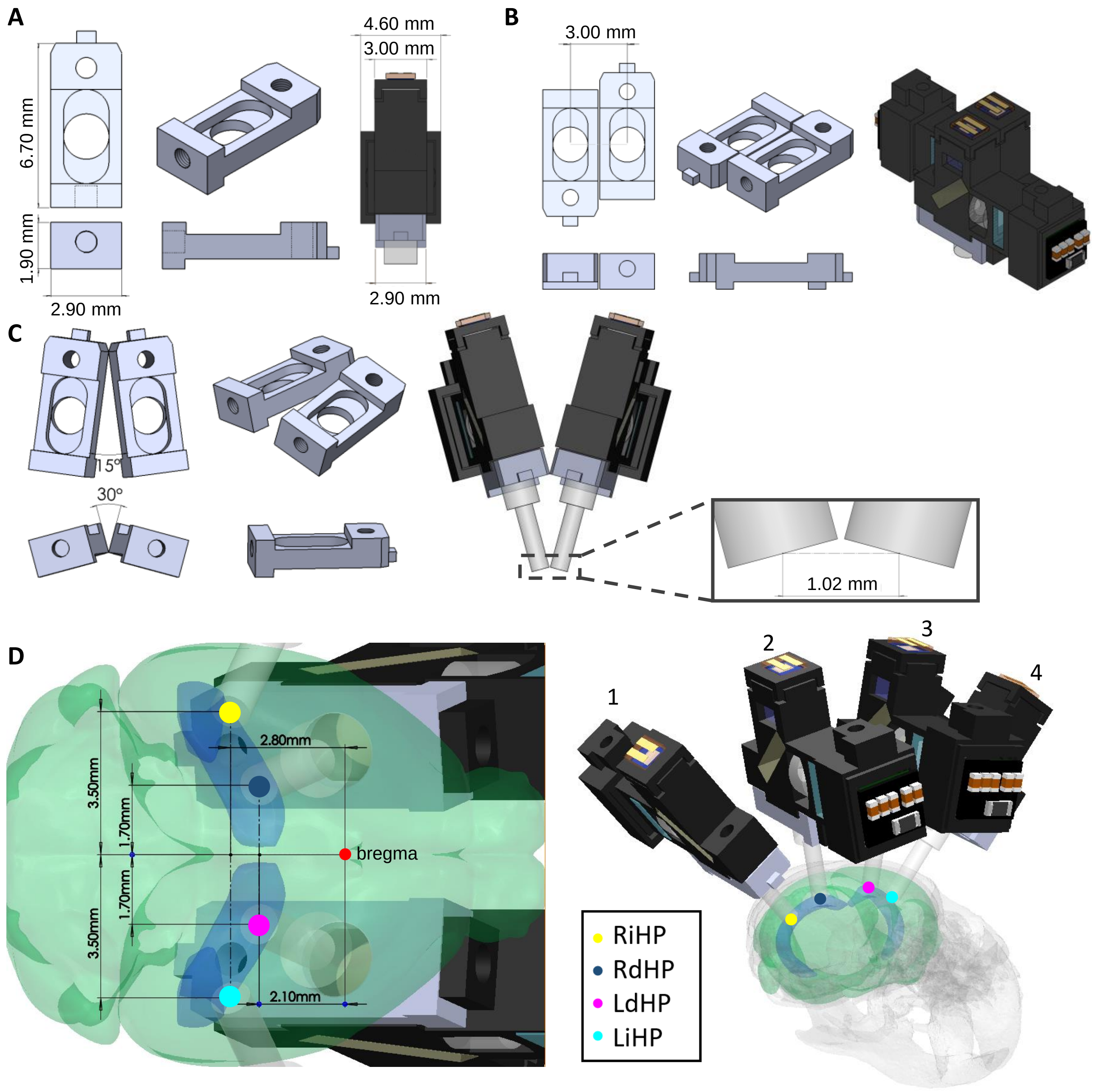


**Supplementary Figure 3. Design of TINIscope arrangement in multiple-region implantations. A,** Engineering drawing of the customized baseplate and an assembled TINIscope with a baseplate (right). Left: top, front, right and isometric views of the baseplate with labeled dimensions. **B,** Two TINIscopes with their baseplates placed side-by-side, resulting in a minimum distance of 3 mm between two GRIN lenses. Left: top, front, right and isometric views of two baseplates placed adjacent to each other; right: 2 TINIscopes assembled with adjacent baseplates. **C,** Tilted arrangement of two TINIscopes achieves a smaller distance between two GRIN lenses (1.02 mm). Left: top, front, right and isometric view of two baseplates placed with a 15-degree angle in the top view and a 30-degree angle in the front view; right, the assembled TINIscopes with the inclined baseplates. **D,** 3D modeling of implanting 4 TINIscopes in the hippocampus from the bottom view (left) and isometric view (right). Blue structure, hippocampal CA1 region; green structure, brain; gray structure, skull.

**Supplementary Figure 4. Design of the commutator system.**

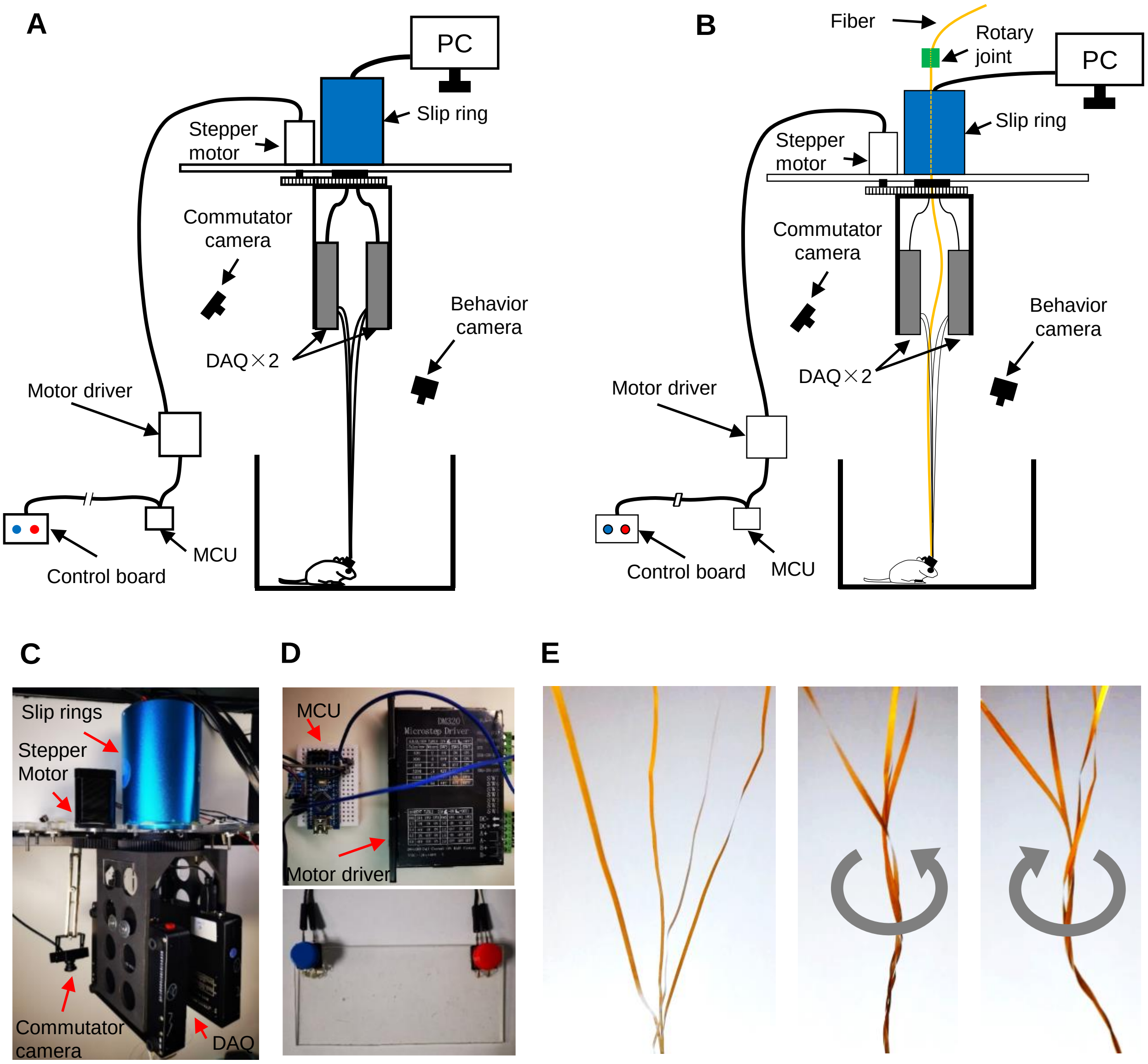


**Supplementary Figure 4. Design of the commutator system. A**, Diagram of the commutator system. DAQ boards were connected to a computer through a slip ring driven by a stepper motor, and the motor was manually controlled by the experimenter outside of the experimental room through the control board. The commutator camera was used for monitoring entanglements of cables, and the behavior camera was used for recording animal behaviors. **B**, Adaptation of the commutator for optogenetic manipulation. The optical fiber was connected to a fiber-optic rotary joint passing through the center hole of the slip ring, and its torque was untangled together with other wires by a stepper motor. **C**, **D**, Photos of the system. Slip ring, motor, camera and DAQ boards in **C**. Microcontroller unit (MCU), motor driver (top) and control board (bottom) in **D**. **E.** Photos of the Flex-PCBs under three different conditions: no twisting, counterclockwise twisting and clockwise twisting.

**Supplementary Figure 5. Comparison of the moving ability of mice carrying no device, 4 TINIscopes, or 4 TINIscope & optical fiber.**

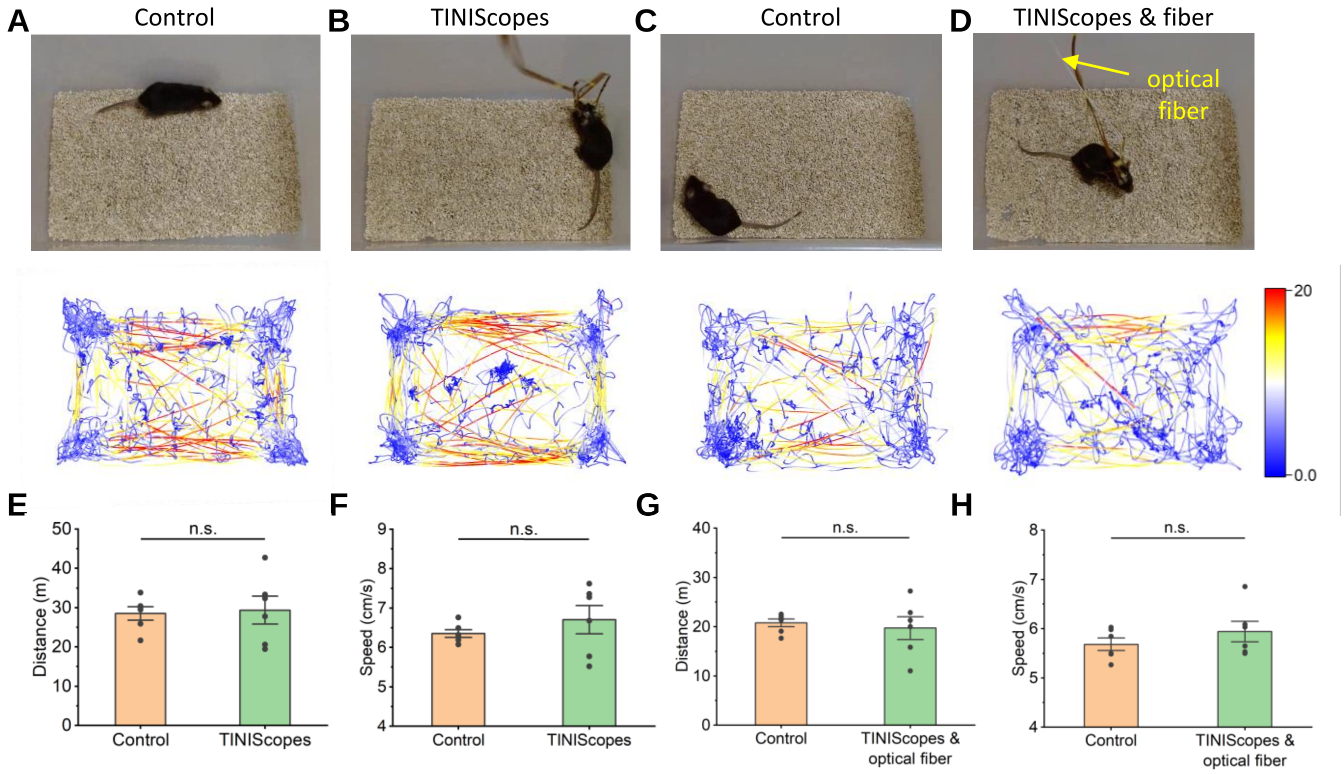


**Supplementary Figure 5. Comparison of the moving ability of mice carrying no device, 4 TINIscopes, or 4 TINIscope & optical fiber. A-D**, Photos of mice in homecage (top) and their trajectories (bottom) in 15-minute trials. **A** and **C** utilized the same group of mice (*n* = 6). Similarly, **B** and **D** utilized the other group of mice (*n* = 6) with 4 TINIscope mounted, while **D** had an additional optical fiber implanted. The comparison pair of **A** and **B** was conducted on day 1, and the comparison pair of **C** and **D** was conducted on day 2. The intertrial interval of each mouse was 24 hours. The color bar indicates the real-time speed (cm/s). **E-F**, Comparison of total distance ($P = 1$) and average speed ($P = 0.47$) between groups **A** and **B**. **G-H**, Comparison of total distance ($P = 0.81$) and average speed ($P = 0.17$) between groups **C** and **D**. n.s., not significant at $p<0.05$ by the Mann-Whitney test.

**Supplementary Figure 6. Longitudinal calcium imaging with TINIscope in the hippocampus.**

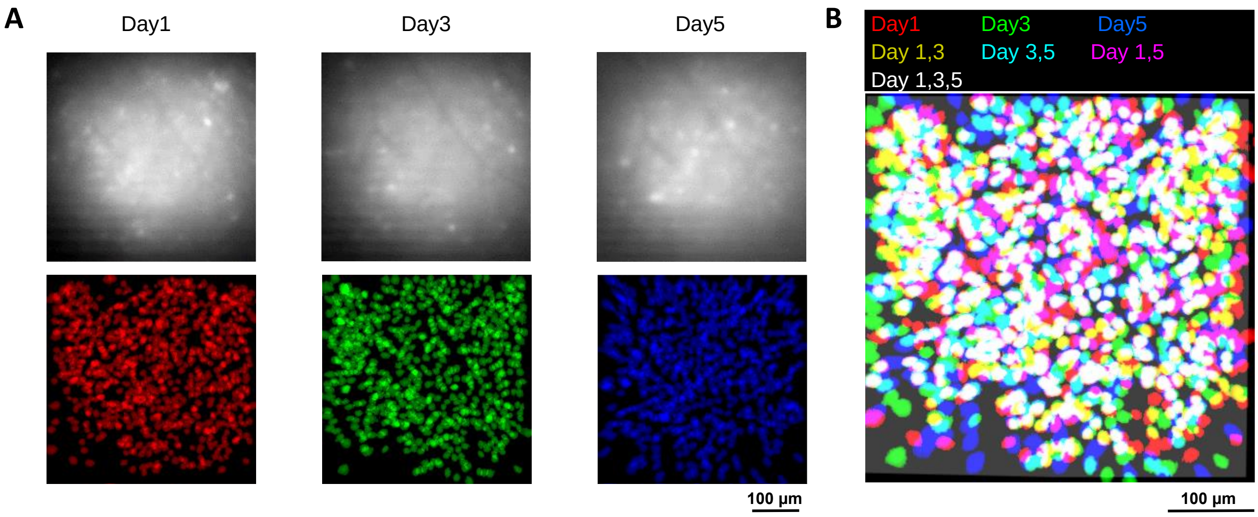


**Supplementary Figure 6. Longitudinal calcium imaging with TINIscope in the hippocampus. A**, Example frames and footprints of the extracted neurons in three different recording sessions. The neuron numbers in three sessions were 677, 614, and 538, respectively. **B**, Overlays of the aligned spatial footprints.

**Supplementary Figure 7. Simultaneous imaging of 4 hippocampal subregions in response to optogenetic ACC stimulation.**

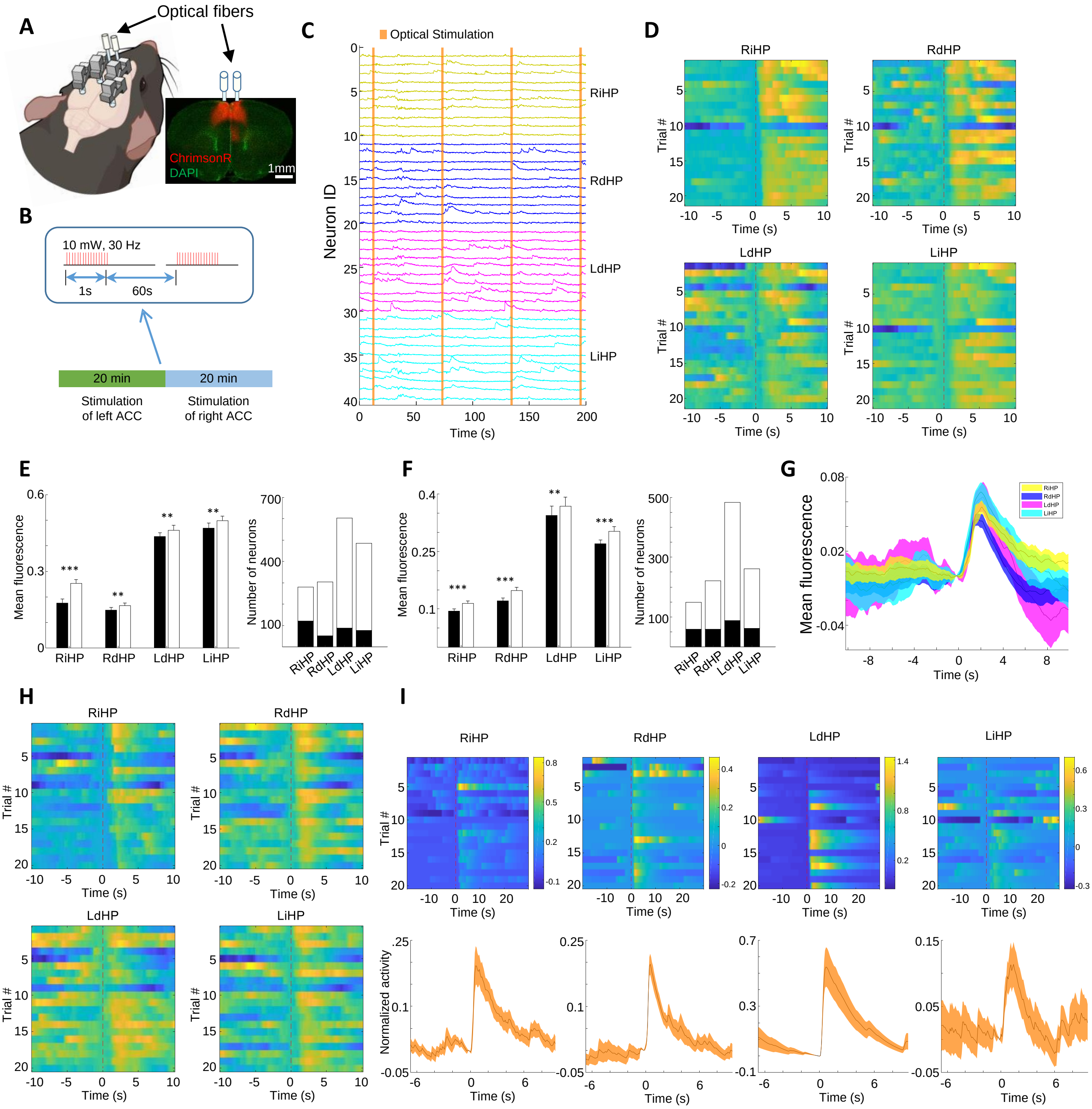


**Supplementary Figure 7. Simultaneous imaging of 4 hippocampal subregions in response to optogenetic ACC stimulation. A**, Left: Illustration of a mouse implanted with 4 TINIscopes in the hippocampus and two optical fibers in the ACC. Right: Fluorescence image of a brain slice showing the expression of ChrimsonR and the locations of optical fiber. **B**, Diagram of the optical stimulation experiments. Each cycle starts with 1 second of 30 Hz stimulation and then rests for 60 seconds. **C**, Example calcium traces of selected neurons. Orange vertical lines indicate the stimulation periods. **D**, Mean population activity aligned with different trials of left-ACC stimulation (t = 0 indicates the stimulation onset), *n* = 21 trials, 1 mouse. **E**, **F**, Left, comparison of the mean fluorescence of all neurons before (black, 2 seconds) and after (white, 2 seconds) ACC stimulation in the left (**E**) or right (**F**). ***$P < 0.001$, **$P < 0.01$, $n$ = 21 trials in **E**, $n$ = 20 trials, 1 mouse in **F**, Wilcoxon matched-pairs sign rank test. The results are presented as the mean±s.e.m. Right, number of neurons that respond (black) vs. those that do not respond (white) to the stimulation. **G**, The mean of the background-subtracted fluorescence signals in 4 regions averaged over all right-ACC stimulation trials ($n$ = 20 trials, 1 mouse). **H**, Same as **D** but with right ACC stimulation. $n$ = 20 trials, 1 mouse. **I**, Example neurons that respond to optogenetic stimulation in the right ACC. Shaded areas in **G** and **I** correspond to the mean±s.e.m.

**Supplementary Figure 8**. **Simultaneous imaging of 4 hippocampal subregions in response to electrical ACC stimulation.**

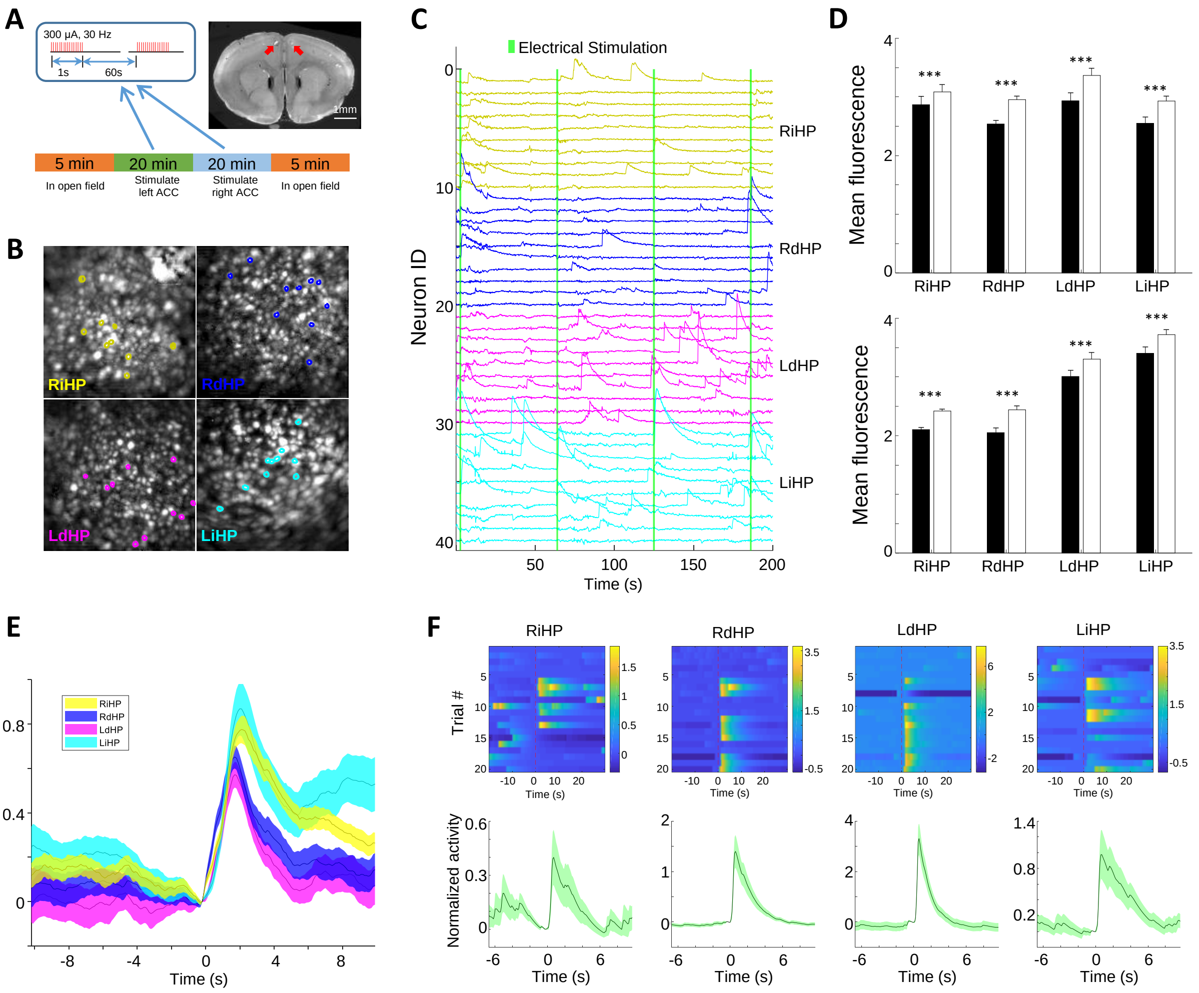


**Supplementary Figure 8. Simultaneous imaging of 4 hippocampal subregions in response to electrical ACC stimulation. A**, Experimental diagram of the electrical stimulations in the ACC, indicated by red arrows in the right panel. Each cycle starts with 1 second of 30 Hz stimulation and then rests for 60 seconds. **B**, MIP of background-subtracted images during electrical stimulation. Contours correspond to example neurons shown in **C**. **D**, Calcium traces of example neurons, and green vertical lines indicate stimulation onsets. **D**, Comparison of the mean fluorescence of all neurons before (black, 2 seconds) and after (white, 2 seconds) ACC stimulation in the left (top, $n$ = 42 trials, 2 mice) or right (bottom, $n$ = 43 trials, 2 mice). ***$P < 0.001$, Wilcoxon matched-pairs sign rank test. The results show the mean ± s.e.m. **E**, The mean of the background-subtracted fluorescence signals in 4 regions averaged over all right-ACC stimulation trials ($n$ = 43 trials, 2 mice). **F**, Example neurons that respond to electrical stimulation in the right ACC. The traces were normalized using the estimated noise level and centered around the value at the onset of stimulation (t=0). Shaded areas in **E** and **F** correspond to the mean ± s.e.m.

**Supplementary Figure 9. Simultaneous LFP recording and calcium imaging in multiple brain regions.**

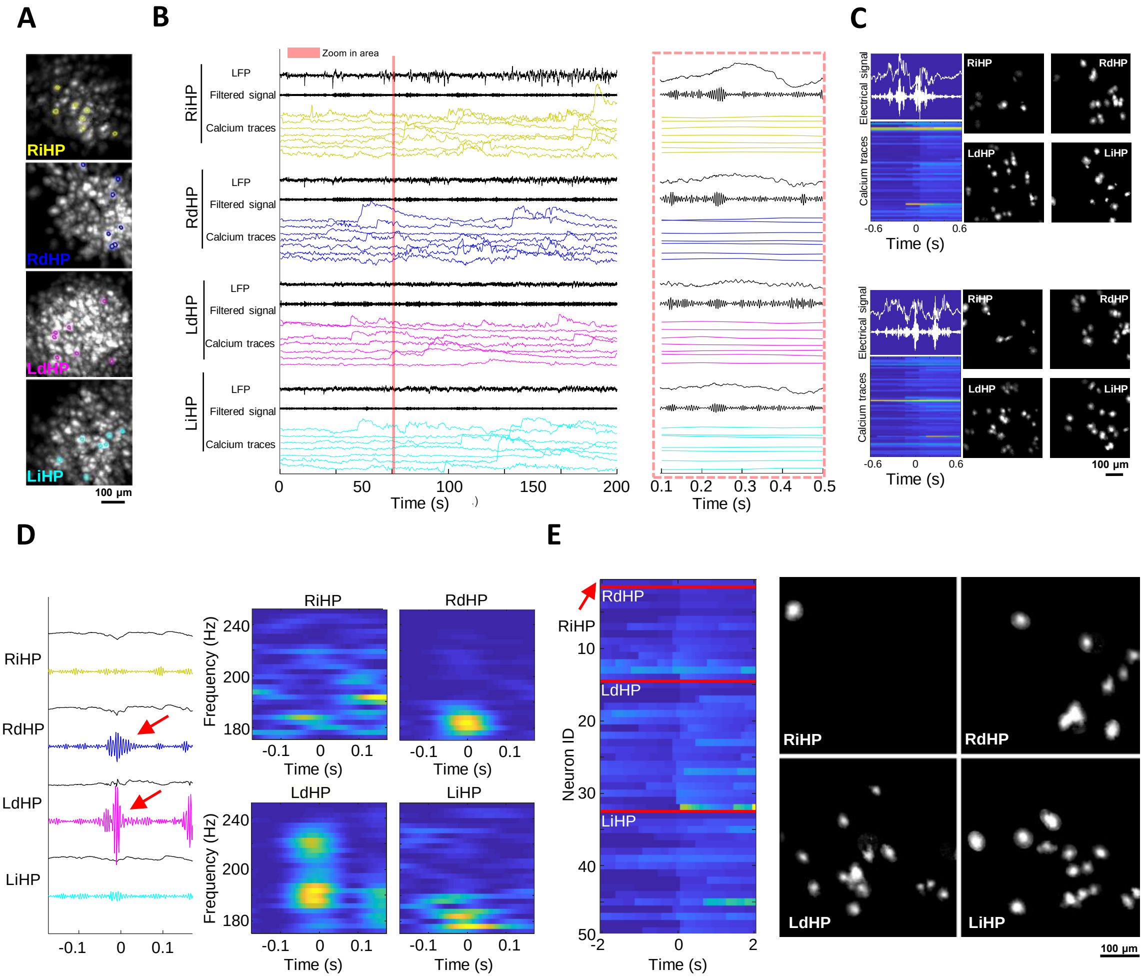


**Supplementary Figure 9. Simultaneous LFP recording and calcium imaging in multiple brain regions. A**, MIP of background-subtracted images. Contours correspond to example neurons shown in **B. B**, Simultaneously recorded LFP signals, bandpass LFP (150-250 Hz), and example calcium traces. The right panel is the zoomed-in view of a 400-millisecond window near the red line in the left panel. **C**, Two example SWRs with synchronous neuronal activity. The spatial footprints of the synchronous neurons are shown on the right. **D**, Raw LFP (black) and filtered signals (150-250 Hz) when 2 regions (red arrows) show SWRs together. Right, spectrograms of LFP signals in 4 regions. **E**, Synchronous calcium traces (left) and their spatial footprints (right) concurrent with the SWR in **D**.

**Supplementary Figure 10. Neuronal assemblies during open-field exploration.**

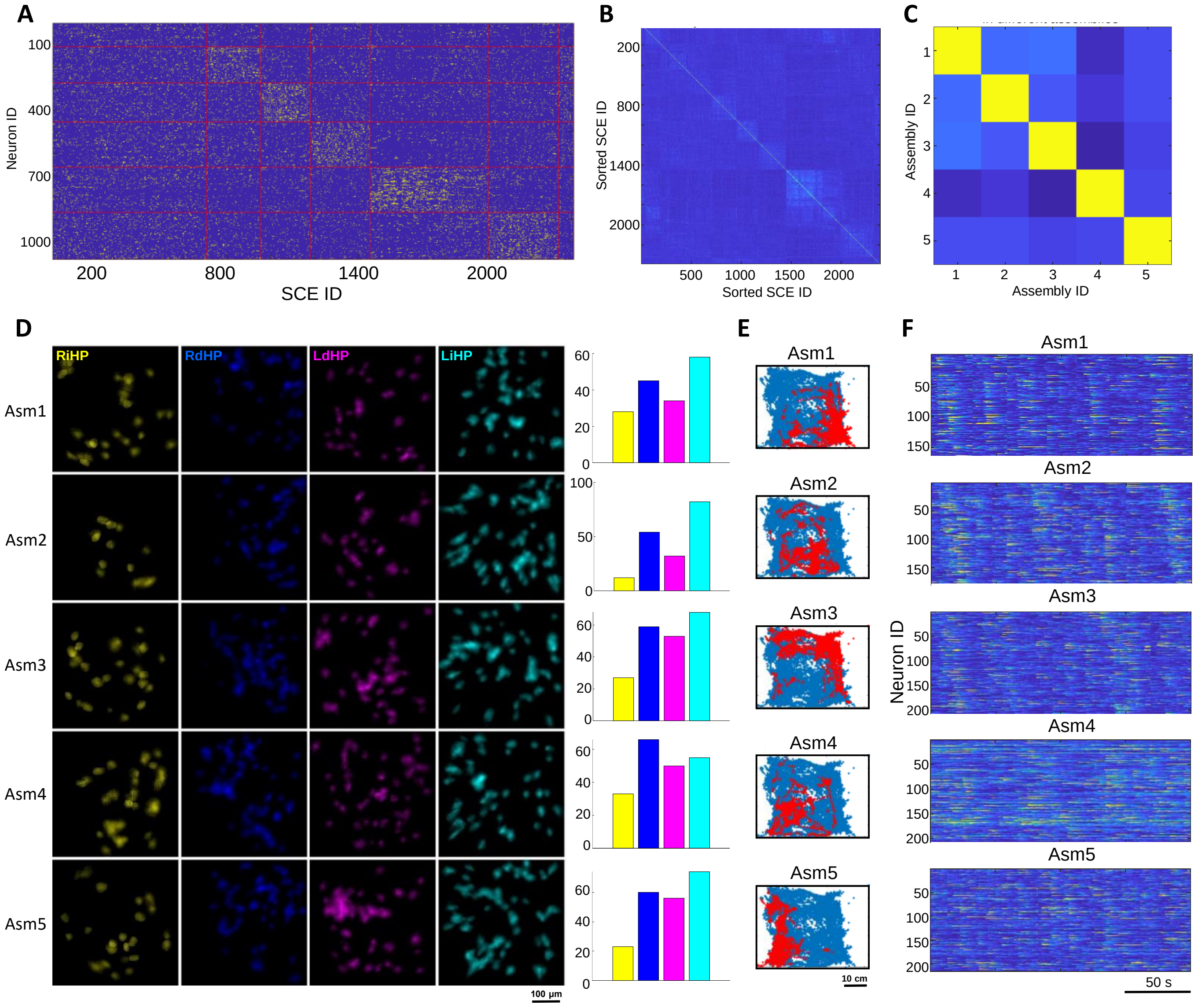


**Supplementary Figure 10. Neuronal assemblies during open-field exploration. A**, Raster plot of all SCEs. Both the neuron IDs and SCE IDs were ordered to match the identified assemblies. **B**, Correlation map of SCEs sorted by their involved assemblies. **C**, Correlation map of all found assemblies. **D**, The neuron number and their spatial footprints in different assemblies. **E**, The mouse's position when SCEs associated with a specific assembly occurred. **F**, Activation sequences of neurons associated with each assembly. A 20-second time window of each SCE (10 seconds before and after the SCE) was selected and then concatenated.

### SUPPLEMENTARY NOTES

##### Supplementary Note 1 Placement of multiple TINIscopes via computer simulation

When implanting multiple TINIscopes, it is essential to design proper equipment arrangements to avoid spatial conflicts and balance weight. To achieve this, Solidworks was utilized for simulating the implantation and determining optimal solutions through three steps. Firstly, a standard 3D model of the whole mouse brain and targeted brain regions was downloaded from the Allen Brain Atlas[1] (https://download.alleninstitute.org/informatics-archive/current-release/mouse_ccf/annotation). Secondly, the downloaded models and 3D model of TINIscopes were imported into Solidworks. Lastly, the GRIN lens of each TINIscope was fixed to the target area, and the orientation of each TINIscope was adjusted to achieve the best arrangement (Supplementary Fig. 3D and Supplementary Video 6). The final orientation of each TINIscope was then exported to guide surgical implantation.

##### Supplementary Note 2 Design of HDI rigid-flex PCB of TINIscope

TINIscope uses a serialized data transmission protocol (MIPI interface) to send CMOS signals to the data acquisition (DAQ) board. This protocol does not require a serializer chip on the head-mounted side, but it needs 10 wires in total for data transmission, SCCB hardware configurations and power supply. Therefore, the conventional coax connection between the CMOS sensor and DAQ board is not suitable for our TINIscope design.

Our solution relies on a customized HDI rigid-flex PCB, which contains a head-mounted rigid PCB and a flexible PCB connected to the DAQ board (Supplementary Fig. 2C, D). The rigid part is a four-layer high-density interconnect (HDI) PCB with blind and buried vias (Fig. 1B, Supplementary Fig. 2C), achieving an almost identical size (3.43 mm × 3.86 mm) to the image sensor. The flex part is a two-layer stack flexible PCB with dimensions of 2.1 mm × 493 mm × 0.19 mm (Supplementary Fig. 2C, D), which is thin, soft and long enough for typical behavior experiments. The DAQ board not only receives CMOS signals from the rigid-flex PCB but also provides the power and clock signals directly, allowing the removal of these modules from the head-mounted side. GUI software was also developed to acquire image data and configure the CMOS image sensors all through HDI rigid-flex PCB.

Flex PCB's susceptibility to electromagnetic interference poses a potential threat to the stability and quality of data transmission. Empirical evidence confirms the feasibility of obtaining reliable CMOS signals through flex PCBs measuring 70 cm or 50 cm in length, as demonstrated by the results. Furthermore, flex PCBs are vulnerable to twisting, bending, and stretching introduced by the movement of mice, potentially impacting data quality. Therefore, our customized commutator becomes indispensable to mitigate these motion-induced effects. Remarkably, with the inclusion of the commutator, we did not encounter any instances of damage to the flex PCBs during our experimental procedures.

##### Supplementary Note 3 Mechanical design of TINIscope

The housing was 3D printed using black resin for the optical and electronic parts. To match the significantly reduced size of the TINIscope main body, the baseplate was adaptively downsized to dimensions of 2.9 mm×6.7 mm×1.9 mm (Supplementary Fig. 3A). When two assembled TINIscopes were placed side by side, they were able to image two regions at least 3 mm apart (Supplementary Fig. 3B). Additionally, the touching areas of the two baseplates were modified to have chamfers, allowing for them to be arranged at a tilted angle. In this way, the minimum distance between the two GRIN lenses was minimized to 1.02 mm. (Supplementary Fig. 3C).

#### MATERIALS AND METHODS

**Optics.** TINIscope's design flips the traditional arrangement of LED excitation and CMOS imaging, placing large CMOS sensors on the side to avoid spatial conflicts when multiple TINIscope devices are implanted. Zemax software was used for optical performance modeling, including the selection and placement of optics. The illumination light from compact LEDs (LMXZ-PB01, Lumileds, 1.3 mm × 1.7 mm) was gathered by a half-ball lens (#45-933, Edmund Optics) and then filtered with an excitation filter (ET470/40x, Chroma) before passing through the dichroic mirror (59002bs, Chroma). Fluorescence emissions were collected using a 1.8 mm diameter GRIN objective lens (#64-537, Edmund Optics) with a 0.52 numerical aperture (NA) and 0 mm working distance. To minimize tissue damage during lens implantation, a 1 mm diameter GRIN relay lens (CLHS100GFT003, Go!Foton) with a 200 μm working distance was bonded to the objective lens using optical adhesive (#37-322, NOA160, Edmund Optics) with a thickness of approximately 100 μm. As a result, the working distance of the objective GRIN lens is shifted by this thickness, which can be compensated by adjusting the slider for image focusing. The emitted fluorescence image was reflected by the same dichroic mirror, focused by a plano-convex lens (#88-775, 3 mm diameter, $f$ = 4.5 mm, Edmund Optics), and then filtered with an emission filter (ET525/50m, Chroma) before being captured by the image sensor. The excitation filter, emission filter, and dichroic mirror were diced to fit the housing size using a dicing saw (ADT7100, AdvancedDicing Tech).

**Image sensor.** The current version of TINIscope verified two image sensor chips from OmniVision (OV6211 and OV7251) with small package dimensions of 3175 μm × 3175 μm/3910 μm × 3410 μm and 8/10-bit RAW output. The chips are equipped with a one-lane MIPI for serial data readout and SCCB for hardware configuration. While the sensor can achieve a maximum frame rate of 120 Hz, it may result in heating issues during long-term recording sessions. Additionally, the Flex-PCBs used may experience greater electromagnetic interference. In practice, the image chips were tailored to collect data at 25/40 fps with 10-bit output in full image size (400 × 400 for OV6211, 640 × 480 for OV7251).

**Electronic circuit design.** Altium Designer was employed to design the electronic circuit. To achieve maximum weight reduction, we relocated most of the CMOS modules, such as the power regulator and clock generator, to the DAQ board (MUD952, Dothinkey). The peripheral circuit adjacent to the image sensor chips retained only the necessary components and was fabricated as a four-layer high-density interconnect (HDI) rigid printed circuit board (PCB) using blind and buried vias. This design allowed us to achieve an almost identical size (3.43 mm × 3.86 mm) to the image sensor. A flexible PCB with a two-layer stack and thickness of 0.19 mm was used to connect the head-mounted PCB with the DAQ board, ensuring unconstrained movement of animals. Each DAQ board handles the MIPI signals from two CMOS sensors and was connected to a personal computer (PC) via a slip ring (B1286-08S-4U1, Senring). The LED uses a flexible PCB and is then connected to the LED driver with a 0.12 mm twisted pair cable.

**Mechanical design.** Solidworks was utilized to design the housing and baseplate of the microscope. The housing was 3D printed using black resin (FLGPBK04, Form 2, Formlabs) at 25 μm axis resolution and contained a main body, a cover for the half-ball lens, and a slider for image focusing. After assembling the half-ball lens, the cover was attached to the main body, and the LED was fixed in place using optical adhesive. The CMOS image sensor chip was also attached to the slider using optical adhesive, and an M0.5 set screw was used to secure the focusing slider when the focal adjustment was done.

The baseplate, made with metal machining, was then bonded to the objective lenses with optical adhesive. To complete the assembly, the housing was secured to the baseplate with two M1 set screws.

**Commutator design.** The commutator was composed of a 3D-printed nylon U-shaped bracket, an electric slip ring (B1286-08S-4U-62641, Senring), a stepper motor (42B60) with a driver (DM320), and a controller (Arduino Nano, Supplementary Fig 4). Two MIPI DAQ boards were mounted on the arms of the bracket, which was fixed to the rotor of the electric slip ring. The stepper motor was coupled to this rotor via a 1:2 gear set, providing the power to rotate the DAQ boards. During experiments, the experimenters determine the rotation direction by monitoring the shapes of flexible PCBs through a camera (Supplementary Figure 4A-C) and then press the controller to rotate the stepper motor clockwise or counterclockwise through the Arduino Nano, avoiding any movement-related wire entanglements.

**Software development.** The application programming interface (API) provided by the MIPI DAQ board was utilized to develop a graphical user interface (GUI) using Qt Creator. Additionally, OpenCV was invoked to handle videos from the behavioral camera. The GUI facilitated the setup of the power supply voltage, main clock frequency, and configuration of the CMOS image sensor. After hardware configuration, all operations relating to the synchronous imaging and data collection from TINIscopes and the behavioral camera were executed through the GUI.

**Measure of optical performances.** The optical resolution of TINIscope is measured by a USAF1951 resolution target. The smallest element of group 7 (linewidth of 2.19 μm) was resolvable, which is decided by the Nyquist limit of the CMOS pixel pitch. In the imaging plane, the full width at half maximum (FWHM) of a 1 μm fluorescence bead was 3.17 μm. To measure the intensity of illumination, we employed an additional microscope to capture images of the illumination light at the focal plane of TINIscope. The radial profile of this image was subsequently analyzed to evaluate the uniformity of the illumination. The focal range of TINIscope is approximately 0 - 200 μm with the adjustment of the slider, measured with a translation stage. The typical LED power was set to 0.05 - 0.35 mW, measured with an optical power meter at the TINIscope focal plane.

**Animals.** Male C57BL/6J mice were housed in a controlled environment with a 12-hour light/dark cycle and ad libitum access to food and water. All animal experiments were conducted following protocols approved by the IACUC (Institutional Animal Care and Use Committee) of Shenzhen Institute of Advanced Technology, Chinese Academy of Sciences (SIAT-IACUC-210802-NS-ZDK-A2028) and the IACUC of the University of Science and Technology of China (USTCACUC192101056).

**Surgery.** Mice, approximately 8 weeks old, were anesthetized with isoflurane vapor (2% for induction and 1% - 1.5% for maintenance) mixed with $O_2$ and fixed on a stereotaxic apparatus. Body temperature was maintained at 37.5 °C using a heating pad during surgery and anesthesia recovery. AAV2/9-hSyn-GCaMP6s virus (Shanghai Taitool Bioscience) was injected into the intermediate hippocampus (iHP) at coordinates of AP: -2.8 mm, ML: ±3.5 mm, DV: -1.4 mm from the surface of the skull with a 35-degree angle and dorsal hippocampus (dHP) at AP: -2.1 mm, ML: ±1.7 mm, DV: -1.4 mm from the surface of the skull with a 9-degree angle. A total of 400 nl of virus was injected at a rate of 15 nl/min per site. After 3 weeks, mice were re-mounted on a stereotaxic apparatus for implantation of the imaging lens. A 1.2 mm-diameter hole was drilled in the skull at each virus injection site, and the cortex above the CA1 at each virus injection site was removed (with a 35-degree angle and

9-degree angle for iHP and dHP, respectively) using a negative pressure pump. Then, a 1 mm-diameter lens was placed on the surface of the CA1 (DV: -1.15 mm below the surface of the skull) and secured to the skull using dental cement. For the optogenetic stimulation experiment, two additional holes were opened above the ACC. A total of 300 nl of AAV2/9-mCaMKIIα-ChrimsonR-tdTomato virus was injected into the ACC (AP: +1.0 mm; ML: ±0.35 mm; DV: -1.5 mm), and a multimode optical fiber with a diameter of 200 μm and NA of 0.37 (Suzhou CooCore Photoelectronic Technology) was implanted above the ACC (AP: +1.0 mm; ML: ±0.35 mm; DV: -1.0 mm). For electrical stimulation, a bipolar platinum-iridium electrode with an interval of 0.2 mm was inserted into the ACC (AP: +1.0 mm; ML: ±0.35 mm; DV: -1.5 mm).

**Histology.** After recording, mice were sacrificed and transcardially perfused with saline followed by 4% paraformaldehyde (Sigma) in PBS. The brains were kept in 4% paraformaldehyde at 4 °C for 24 hours and subsequently immersed in 30% sucrose for 72 hours before being sliced into 40-μm coronal sections and imaged under a fluorescence microscope (Olympus MVX10).

**Combination of TINIscope and optogenetics.** The ACC neurons expressing ChrimsonR were stimulated with a 593 nm laser (Shanghai Fiblaser Technology) via an optical fiber attached to the commutator system to avoid entanglements. The fiber was connected with a fiber-optic rotary joint placed on top of the slip ring. The excitation filters and LEDs were replaced with ET440/40x (Chroma) and LXZ1-PR01 (Lumileds), respectively, to minimize potential crosstalk between the excitation of GCaMP6s and ChrimsonR. An LED is positioned on the behavioral cage, receiving a shared trigger signal with the BNC connector of the laser. This ensures that the onset of stimulation aligns with the activation of the LED, enabling direct detection of this synchronization from the behavior videos. Therefore, we can effectively synchronize optogenetics with behavior using this setup, and a similar synchronization strategy was used for electrophysiological stimulation and recordings.

**Combination of TINIscope and electrophysiology recordings.** A single platinum-iridium electrode was affixed to the side of each imaging lens, with the tip of the electrode positioned 0.3 mm above the lens surface. The electrode was then implanted into the CA1 along the imaging lens. Two additional ground wires connected to the skull screw were placed near the bregma and lambda, respectively. The recording electrodes and ground wires were connected to an electrical socket. Then, the lens, socket and screws were secured to the skull using dental cement. A digital head stage for analog-to-digital conversion of neuronal signals was plugged into the socket. The signals were sampled at 30 kHz and transmitted to the data acquisition equipment (NeuroStudio System, Jiangsu Brain Medical Technology Co.). The electrical cables were connected to slip rings to avoid entanglement with the TINIscope system. Additionally, a shielding mesh (100 cm × 80 cm × 120 cm) covered the whole system to minimize interference with electrophysiological signals.

**Evaluation of mouse mobility**. A total of 12 male C57BL6/J mice were divided into two groups: control ($n$ = 6) and TINIscope-carrying (TC, $n$ = 6) groups. For the TC group, the same surgical procedures described earlier were performed to implant the GRIN lens and mount 4 TINIscopes. The control mice remained unaltered. Two separate experiments were conducted on consecutive days. On day 1, each mouse in both groups was allowed to explore a homecage with new padding for 15 minutes per trial. All cables were connected to the TC mice to simulate actual recording sessions. On day 2, an additional optical fiber was added to the TC group, and the same experiments as on day 1 were repeated. Since mice were less likely to explore the same homecage on day 2, we assessed their performance separately for each day (see

Supplementary Figure 5). The body centers of mice were extracted using DeepLabCut, followed by calculating the total distances and the average speeds with custom MATLAB scripts. Due to the potential obstruction of mouse body parts by TINIscope devices or wires, DeepLabCut may encounter difficulties in recognizing their precise locations or lead to missing specific markers. To mitigate this problem, we manually inspected the result and labeled the outliers, which were further fixed by utilizing the refining-outlier function in DeepLabCut.

**Behavioral experiments.** Our behavioral experiments included the T-maze and context-changing open-field tests. Both boxes were made of polymethyl methacrylate (PMMA). The T-maze box had dimensions of 40 cm × 44.5 cm × 20 cm, with a vertical arm length of 34.5 cm, a horizontal arm length of 40 cm, and an arm width of 10 cm. The top of the vertical arm was designated as the starting area, while the two ends of the horizontal arm were designated as the reward zones. The starting area was equipped with an infrared sensor to detect the presence of the mouse, as well as two LEDs that signaled the start of a trial and whether a reward was received. In addition, the wall at each end of the maze contained a 1 cm hole with an infrared sensor to detect nose poke behavior, and a metal pipe was used to deliver water. The extra LEDs on each side of the horizontal arm indicated nose poke behavior. Sensors and LEDs were connected to an Arduino Nano, which utilized a pre-uploaded program to control the start and end of each trial, deliver water through a peristaltic pump, and transmit results to a computer. Bonsai[2] software was used to receive information from the Arduino Nano. Prior to the test, mice were deprived of water for 24 hours. At the start of each trial, the mouse was placed in the starting area, the LED indicating the start of the trial was illuminated, and the program in the Arduino Nano randomly selected one of the two end holes as the reward point. The mouse was allowed to explore the entire T-maze, and the trial ended only when the animal poked the reward point and received 15 µl of water. The mouse was required to return to the starting area to initiate the next trial. The test concluded once the mouse had completed 20 trials or after a duration of 1 hour. The context-changing open field test was conducted in a box with dimensions of 30 cm × 30 cm × 20 cm. The test consisted of four stages. First, the mouse was placed in the box for 10 minutes to explore freely. Next, three platforms of different heights (3 cm, 6 cm, and 9 cm) were added to divide the box into four areas with varying heights, and the mouse was allowed to explore for 15 minutes. In the third stage, a small piece of chocolate was placed at the center of the highest platform, and the mouse was allowed to explore for an additional 15 minutes. In the final stage, all platforms were removed, and the mouse was allowed to explore the original open field for 10 minutes. The locations of the mice at each time point were identified from behavioral video using Bonsai and then converted into centimeters.

**Calcium data analysis.** Raw video data were analyzed using NoRMCorre for nonrigid motion correction and CNMF-e[3] for neuronal trace extraction. Spatial (2x) and temporal (4x) down-sampling was usually used to yield a faster processing speed without noticeable information loss. The patch size in nonrigid motion correction was 80 × 80 pixels. After the first round of CNMF-e, the extracted neurons with peak-to-noise ratio (PNR) values less than 5 were identified as false positives and deleted. The remaining neurons were then manually screened depending on their shapes and traces, followed by an additional round of temporal trace refinement. CellReg software was utilized to register neurons across multiple sessions[4].

**Spatially modulated cell identification.** Spatially modulated cells were identified by computing how much spatial information $I$ was provided by neuronal activity as previously reported[5-7] through custom MATLAB functions. Locations were 40 × 40 pixel binned, and only bins with occupancies longer than 0.5 seconds were considered. The neuronal activity value was the spatially Gaussian-smoothed ($\sigma$ of 0.5 bin) spiking activity, referred to as deconvolved traces in the CNMF-e results, when the speed of the mouse exceeded 30 pixels/s (~0.8 cm/s). Data For each neuron, $I$ was computed as:

$$I = \sum_i p_i a_i \log_2(a_i/\bar{a})$$

where $i$ is the location bin number, $p_i$ is the probability that the mouse was in bin $i$, $a_i$ is the average neuronal activity in bin $i$, and $\bar{a}$ is the average neuronal activity through all bins. If $I$ exceeded the 95th percentile of 100 shuffled $I$, the cell was identified as a spatially modulated cell. To generate each shuffle, the time course was circularly shifted by a minimum of 500 frames, divided into six chunks, and their order was permuted. After identifying a neuron as a spatially modulated cell, its place fields were represented by plotting the $a_i$ of all bins.

**Decoding analysis.** Decoding mouse locations from hippocampal neuronal activity was performed with the neural decoding package [8]. The LSTM algorithm with specific hyperparameters (units = 200, dropout = 0.25, number of training epochs = 10) was selected because it has empirically higher decoding accuracy. The decoder generates the x and y positions of the mouse in one temporal bin (bin size = 100 ms) using the population neurons' calcium traces in the previous four temporal bins along with the current bin as input. Backpropagation was utilized to train the LSTM, minimizing the discrepancy between the network's predictions and the true mouse positions. Prior to decoding, the extracted calcium traces were temporally aligned with mouse locations, and silent periods at the start or end of trials were manually removed. The resting data were split into 10 folds equally, with 9 folds used for training an LSTM decoder and the last fold used for calculating prediction error, using a 10-fold cross-validation procedure. To determine the chance level of decoding error, the same procedure was performed on location-shuffled data, where the location vector was flipped in time and randomly cycle-shifted for at least 2000 frames.

**Stimulation modulated cells.** All extracted neuronal activities were first aligned with the onset of each stimulation, and then the mean activities 2 seconds before and after the stimulation were compared. Neurons showing significantly higher or lower ($P < 0.05$, Wilcoxon matched-pairs signed rank test) activity after stimulation were identified as stimulation-modulated cells. The population activity in each region was averaged from the extracted fluorescence signals of all neurons in that region.

**SWR detection.** Extracellular signals recorded from the NeuroStudio system were down-sampled to 1000 Hz, filtered at 150-250 Hz with a zero-phase digital filter using a 256th-order Hamming window, and subsequently z-scored. Events with peak values higher than 7 standard deviations (SDs) and 20 ms durations with peak values higher than 3 SDs were detected as SWR events.

**Statistics.** All statistical analyses were performed with Python and MATLAB. The Mann-Whitney test, Wilcoxon matched-pairs signed rank test and paired t-test were used for statistical analysis.

**Code and design availability.** All the code used and hardware design can be found on our open-source GitHub repositories (https://github.com/TINIscope/TINIscope). Within this repository, there is a comprehensive list containing purchase links and prices for all components needed. Additionally, we have provided a detailed step-by-step assembly guide, which requires some expertise in electrical and optical engineering.

### CAPTIONS FOR SUPPLEMENTARY VIDEO

**Supplementary video 1**

Example video data of a single-TINIscope recording in the mouse hippocampus. The neurons were infected with AAV2/9-hSyn-GCaMP6s. The video is a 20x speedup.

**Supplementary video 2**

Concurrent 4-region calcium imaging and behavioral recording of freely moving mice in open-field (section 1) and T-maze (section 2) environments. Section 1 is an open-field test where the mouse climbed to the highest platform for fetching chocolates, and Section 2 is a T-maze test where the mouse was exploring to find a water reward and then moved to the starting point. The videos in the middle show the raw and background-subtracted calcium imaging in 4 hippocampal subregions (RiHP, RdHP, LdHP, LiHP). The right shows calcium traces of example neurons. The video is 4x speedup.

**Supplementary video 3**

Simultaneous electrical stimulation and 4-region calcium imaging in the open-field experiment. The videos in the middle show the raw and background-subtracted calcium imaging in 4 hippocampal subregions (RiHP, RdHP, LdHP, LiHP). The right panel shows calcium traces of example neurons, and green lines indicate the electrical stimulation. The video is 4x speedup.

**Supplementary video 4**

Simultaneous electrophysiological recording and multi-region calcium imaging. The videos in the middle show the raw and background-subtracted calcium imaging in 4 hippocampal subregions (RiHP, RdHP, LdHP, LiHP). The right panel shows the concurrently recorded LFP signals, filtered signals (150-250 Hz) and calcium traces of example neurons. The video is 4x speedup.

**Supplementary video 5**

Decoding accuracy of mouse positions from extracted neuronal activity in open-field (section 1) and T-maze (section 2) environments. Green circles indicate the true position identified by Bonsai, while the others indicate the predicted mouse positions from the extracted population activity of all regions (red dots), RiHP (yellow crosses), RdHP (blue crosses), LdHP (magenta crosses), and LiHP (cyan crosses). Note that decoding with all 4 regions yields superior performance to decoding with any single region. The video is 4x speedup.

**Supplementary video 6**

Computer simulation of implanting multiple TINIscopes in a single mouse brain. TIINIscopes were first placed at targeted brain sites, and then their orientations were adjusted to achieve collision avoidance and weight balancing (section 2).
